# Scalable Image-Based Quantification of Cell Permeability and Actin Remodeling: An Example in a Gut-on-Chip Platform

**DOI:** 10.1101/2025.04.04.647153

**Authors:** Moran Morelli, Karla Queiroz

## Abstract

Evaluating cellular responses to toxic compounds is essential for assessing the safety and potential hazards of drugs, environmental pollutants, and food contaminants. Traditional in vitro methods often lack the precision and scalability required for comprehensive toxicological assessments. This protocol presents an efficient and scalable approach for quantifying cellular damage and toxicity using organ-on-chip technology, specifically the OrganoPlate® platform. By combining fluorescent probes—DRAQ7 for cell membrane integrity, ActinGreen for cytoskeletal changes, and NucBlue for nuclear counting—with high-throughput image analysis via CellProfiler, this method provides detailed and quantitative insights into cellular damage induced by toxic compounds. The protocol includes step-by-step instructions for staining, image acquisition, and data analysis, as well as troubleshooting guidance. CellProfiler’s open-source nature, flexibility, and automation capabilities enable reproducible, high-throughput workflows, offering significant advantages over traditional manual image analysis. Its ability to assess cytotoxicity in human tissue models makes this protocol a valuable tool for safety testing in drug discovery and environmental toxicology. Furthermore, the approach is highly adaptable, accommodating a variety of cell types and toxic compounds, and is well-suited for rapid screening and risk assessment across diverse research and industrial applications.

## Introduction

Toxin exposure is a widespread issue that poses significant risks to human health. Environmental contaminants, pharmaceuticals, and food-borne agents are common sources of chemical exposure that can potentially harm the human body [1]. One of the primary sites of exposure to these chemicals is the gastrointestinal tract, which plays a crucial role in protecting the body against these substances [2]. Understanding cellular responses to toxins is essential for assessing the risks posed by environmental and biological agents. This approach can also be extended to drug safety testing, where evaluating the cytotoxicity of therapeutic compounds is vital during drug development.

Cell-based screening methods provide cost-effective, ethical alternatives to animal models for evaluating chemical safety and efficacy[3]. These methods are especially useful in studying the effects of chemicals on intestinal models, offering insights into how substances are absorbed, metabolized, and excreted in the gut [4]. However, traditional *in vitro* models often fail to predict toxicity in human accurately [5].

To address this, advanced systems like the OrganoPlate platform have been developed. The OrganoPlate is a membrane-free, high-throughput organ-on-a-chip platform that allows cells to be cultured in direct contact with the extracellular matrix (ECM) and under continuous media perfusion. This platform supports large-scale experiments across 64 microfluidic chips, and more accurately mimics intestinal differentiation, polarization, and gene expression [6]. It has been successfully applied in studies on intestinal inflammation [7], drug permeability [8], and toxicology [9–11].

This protocol describes the quantification of actin remodeling and DRAQ7-permeability in OrganoReady® Colon Caco-2 (Figure 1). Quantifying cellular responses to toxins is critical, with the actin cytoskeleton being a key target for many toxins. Actin remodeling, such as depolymerization or abnormal reorganization of filaments, can disrupt cell shape, signaling, and barrier integrity, potentially leading to cell death [12, 13]. Measuring these changes is an early marker for cellular damage. Similarly, DRAQ7, a fluorescent dye that penetrates only damaged cells, is another useful metric for evaluating cytotoxicity. DRAQ7 staining helps distinguish between live and dead cells, providing precise quantification of membrane damage caused by toxic stress.

**Figure 1.**
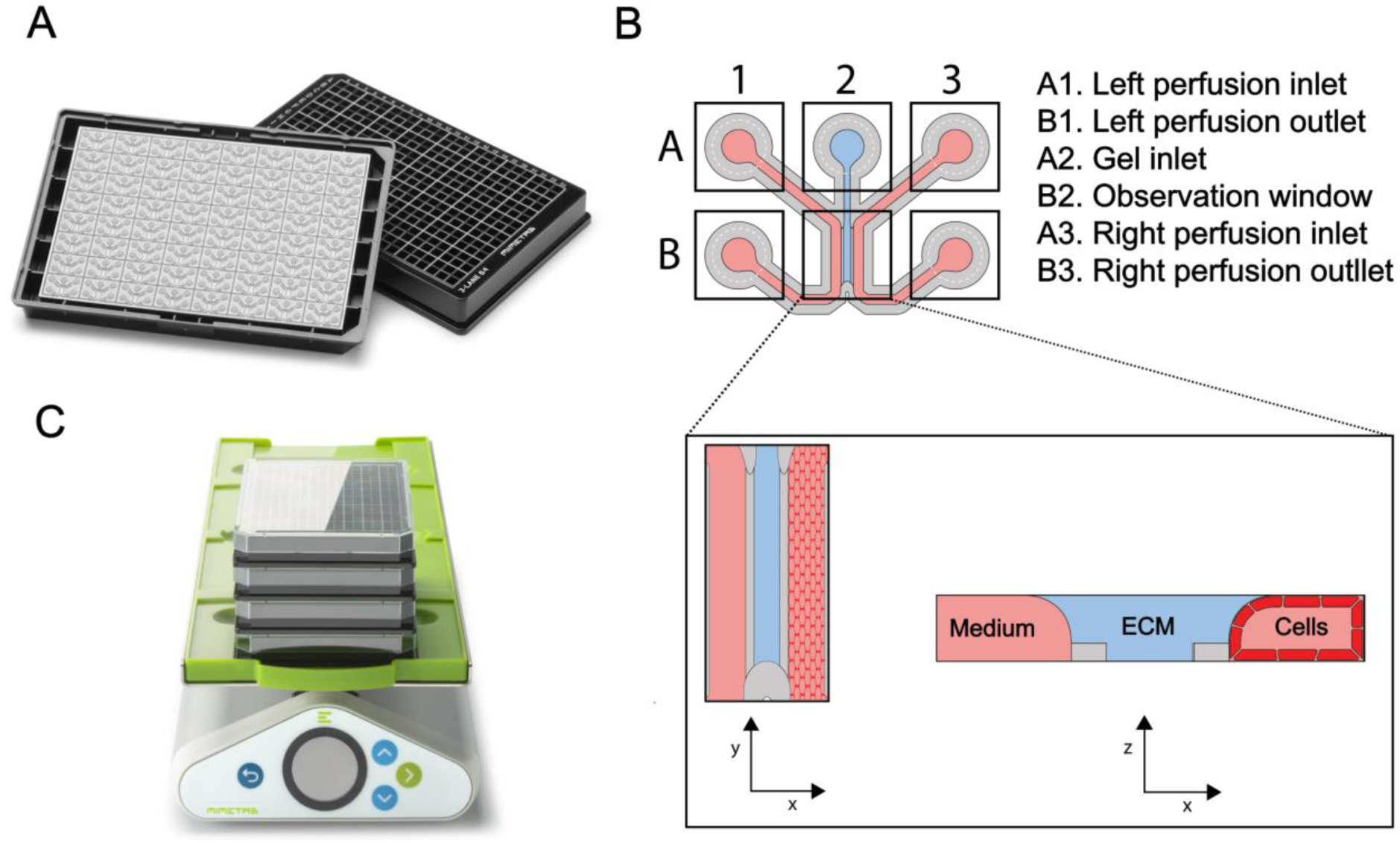
The OrganoPlate technology. (**A)** The OrganoReady® Colon Caco-2 platform, based on the OrganoPlate® technology, consists of 64 microfluidic chips embedded in a standard 384-well microtiter plate. The chips come pre-seeded with Caco-2 tubules, allowing for immediate experimental use without the need for cell culture and differentiation. (**B)** Schematic of a single microfluidic chip, which consists of three channels: the left perfusion channel accessible via inlets A1 and outlet B1; the middle gel channel accessible through inlet A2; and the right perfusion channel accessible through inlet A3 and outlet B3. Squares represent the access wells of the 384-well plate. Imaging and direct observation of the culture is possible via the observation window B2. Top and cross section are shown in the box, with the caco-2 tubule on the right perfusion channel, the middle channel seeded with extracellular matrix (ECM) and the left channel filled with medium. **(C)** Illustration of the OrganoFlow rocker, on top of which OrganoPlates are placed to induce medium flow.

Here, we present a streamlined, reproducible method for assessing cytotoxicity and cellular damage in intestinal tubules using the OrganoPlate platform. This approach leverages automated image analysis through CellProfiler to provide quantitative, high-throughput data on cellular damage, which enhances both the reproducibility and scalability of cytotoxicity assessments. Unlike traditional in vitro models, which often lack predictive accuracy for human toxicity, the OrganoPlate offers an improved representation of human tissue environments, making it a valuable tool for studying chemical exposure in intestinal models.

### Staining protocol

Actin and DRAQ7 staining in the OrganoPlate have been described in literature [10, 11]. Here we used the same protocol, where DRAQ7 is added before fixation, plates are fixated and then actin and nuclei staining are performed.

1. Add the DRAQ7 fluorescent probe before fixation:
  a. Prepare DRAQ7 solution by diluting the stock vial 1:100 in cell culture medium or HBSS. 50uL of solution is needed per chip, so we suggest preparing 60uL/chip to account for pipetting error.
  b. Aspirate all wells of the OrganoPlate
  c. Add 25uL of DRAQ7 staining solution to the tubule inlet, and 25uL in the tubule outlet.
  e. Place the plate back on the OrganoFlow rocker within the incubator for 30 minutes
  f. After incubation aspirate the staining solution and proceed to fixation of the plate
2. Bring the plate to a fume hood to perform fixation:
  a. Prepare fixation solution by diluting formaldehyde to 3.7% in HBSS with calcium and magnesium. 300uL/chip are needed, so we recommend preparing 360uL/chip to account for pipetting error.
  b. Aspirate the medium in all inlets/outlets of the OrganoPlate
  c. Add fixation solution to all tubules inlets and outlets and incubate for 15 minutes
  d. After incubation, aspirate the fixation solution and wash by adding PBS to all tubule inlets/outlets following the same pipetting scheme as for the fixation solution. **Note**: fixated plates can be stored for up to 2 weeks at 4 degrees. For storage, seal the plate with an adhesive film and seal the edges with parafilm. When staining a plate that was stored at 4 degrees, let the plate warm up to room temperature before proceeding with the staining.
3. ActinGreen and NucBlue staining
  a. Prepare permeabilization solution (0.3% Triton X-100 in PBS)
  b. Remove the PBS in all inlets/outlets of the OrganoPlate
  c. Add permeabilization solution in all tubules inlets and outlets and incubate for 10 minutes, static at room temperature, following the same pipetting scheme as for the fixation solution.
  d. While the plate is incubating, prepare staining solution by adding 2 drops/mL of ActinGreen and 2 drops/ml of NucBlue in PBS. 50uL of solution is needed per chip, so we suggest preparing 60uL/chip to account for pipetting error.
  e. After incubation, wash the plate 2 times with PBS
  f. Add 25uL of staining solution to the tubule inlet, and 25uL in the tubule outlet.
  g. Place the plate on rocker at room temperature and incubate for 30 minutes
  h. Wash chips with PBS
  i. Aspirate all wells (also observation windows and gel in/outlets)
  j. Add 50uL PBS to all wells and image the plate.

**Note**: stained plates can be stored at 4 degrees for up to a week, although we recommend imaging within 2 days after staining. For storage, seal the plate with an adhesive film, seal the edges with parafilm, and wrap the plate in aluminum foil.

**Tip:** when pipetting the same volume in the inlet and outlet of the same channel, always make sure to start with one side first and then add to the other side. If added at the same time, the chance of creating a bubble is higher.

### Data acquisition

Fluorescent images are acquired with a Micro XLS-C High Content Imaging Systems (Molecular Devices) using a 10X objective and taking max projections of the bottom of the tubule.

**Note:** Any high content imager can be used.

1. Select the observation window wells of the chips to be imaged
2. Configure the imager to capture three wavelengths:

a. NucBlue: excitation at 360 nm and emission at 460 nm
b. ActinGreen: excitation at 495 and emission at 518 nm
c. DRAQ7: excitation at 599/644 nm and emission at 697 nm
3. Perform alignment and focusing tasks for each chip to guarantee the quality of the images captured.

### Quantification protocols

The purpose of this step is to quantify the images obtained from the fluorescent probe staining. It is performed in the open-source software CellProfiler 4.2.6. where data are loaded, sorted and objects are identified and quantified in all wavelengths.

To download open-source software CellProfiler, use this link: https://cellprofiler.org/releases

#### CellProfiler modules

CellProfiler operates through a series of modular workflows, where each module performs specific tasks in image processing, object identification, and data analysis. Here are the key modules used in this protocol:

**Table 1.** CellProfiler modules used in this protocol.

| Module Name | Description | Used For | Category |
| --- | --- | --- | --- |
| Images | Loads and manages images for analysis. | Importing fluorescence images from Micro XLS-C System. | Input/Output |
| Metadata | Extracts metadata (information about images/experiments) from filenames/folders. | Extracting experiment name, well positions, wavelength. | Input/Output |
| NamesAndTypes | Assigns names to images based on matching rules (e.g., wavelengths). | Renaming image channels to specific probes (e.g., NucBlue, ActinGreen, DRAQ7). | Input/Output |
| CalculateMath | Performs mathematical operations on measurements extracted from objects. | Calculating % of DRAQ7-positive nuclei to assess cell membrane integrity. | Objects Processing |
| ConvertObjectsToImage | Converts object outlines or masks into binary or grayscale images, representing identified objects as pixel intensities. | Saving object masks | Image Processing |
| Crop | Crops images to focus on specific regions of interest. | Isolating microfluidic chip regions containing cells. | Image Processing |
| ExportToSpreadsheet | Exports quantitative measurements and metadata associated with identified objects or images into spreadsheet formats (e.g., CSV files). | Export measurements into a spreadsheet for analysis. | File Processing |
| IdentifyPrimaryObjects | Identifies primary objects (e.g., nuclei or cells) based on intensity/shape criteria. | Detecting and quantifying Hoechst/DRAQ7-positive nuclei and Actin-positive objects. | Objects Processing |
| FlipAndRotate | Rotates or flips images based on specified angles or directions, adjusting image orientation for consistency or specific analysis requirements. | Rotating images to have the tubules horizontally when working with 3-lane 64 OrganoPlate format | Image Processing |
| MeasureImageAreaOccupied | Measures the area occupied by objects in the image. | Quantify the are occupied by actin clumps | Measurement |
| OverlayOutlines | Generates overlays that visually annotate identified objects (e.g., nuclei, cells) on microscopy images. | Used to visualize and verify object identification accuracy before proceeding with further analysis steps. | Image Processing |
| RelateObjects | Links or relates one set of identified objects with another. | Link Hoechst-positive and DRAQ7-positive nuclei to quantify double-positive objects. | Objects Processing |
| SaveImages | Saves processed or annotated images for documentation or further analysis. | Storing images showing identified objects and measurements. Particularly useful to visualize and verify object identification before proceeding with further analysis steps. | File Processing |

**Note:** The “Input/Output” category in CellProfiler consists of modules that manage data flow into and out of the software. Unlike other modules that perform specific image processing or object analysis tasks, Input/Output modules are integral and always present by default. They handle tasks such as importing images, extracting metadata, and organizing data files, ensuring efficient integration and management of experimental datasets within CellProfiler’s analytical framework.

#### Image pre-processing

##### loading and sorting images

1. Open CellProfiler and load the images in the designated section of the “Images” module (**Figure 5. CellProfiler image loading interface.** Go to image module and right click on the box highlighted in red and load images to be analyzed (drag and drop is also possible).**Figure 5**):
  a. Ensure “Images” module is selected
  b. Go to the folder where the images are stored
  c. Select the images and drag and drop them in the specified field **Tip:** when starting a new protocol, go to CellProfiler preferences (file è preferences) to set the default input and output folders. This will ensure the images are taken from and saved in the right folder. **Note:** We recommend loading only a few images (including controls) when designing the pipelines. Once the pipeline is ready, the whole dataset can be loaded and analyzed.
2. Use the “Metadata” module to extract information from the image file or folder name.
  d. Select the “Metadata” module
  e. Go to “Extract Metadata” and click “Yes”
  f. Set “Metadata Source” to “filename”
  g. Use the regular expression editor to extract the following information:
    - Experiment name
    - Well
    - Wavelength
  h. Check the output table to make sure the extracted information is correct
3. Use the “NamesAndTypes” module to assign a name to images. This allows to rename the wavelength with the probe name.
  i. Select the “namesAndTypes” module
  j. Go to “Assign a name to” and select “Image matching rules”
  k. Use the selection criteria to extract the wavelength from the image filename and rename it (this operation needs to be repeated for each wavelength).
  l. Check the output table to make sure the extracted information is correct

**Note:** wavelength information can also be extracted from the metadata

**Figure 2.**
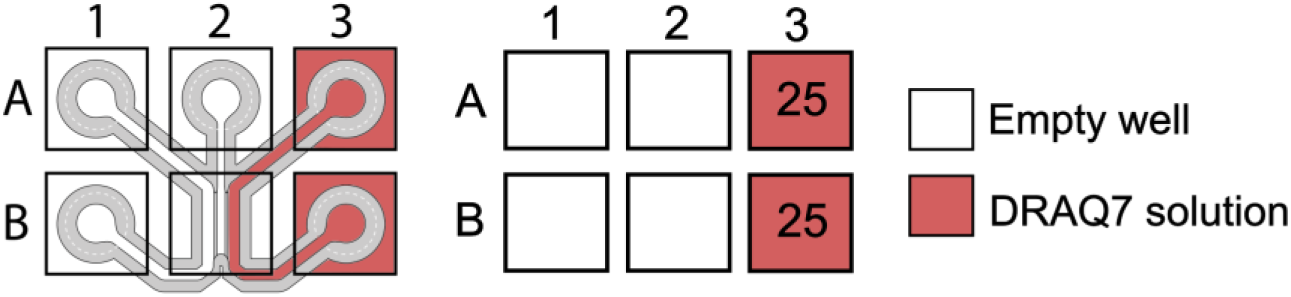
Pipetting scheme for DRAQ7 staini**ng**.

**Figure 3.**
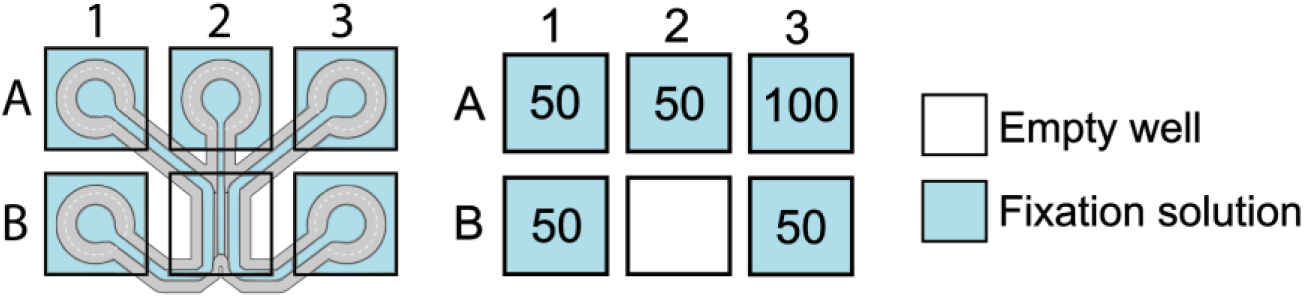
Pipetting scheme for fixation solution.

**Figure 4.**
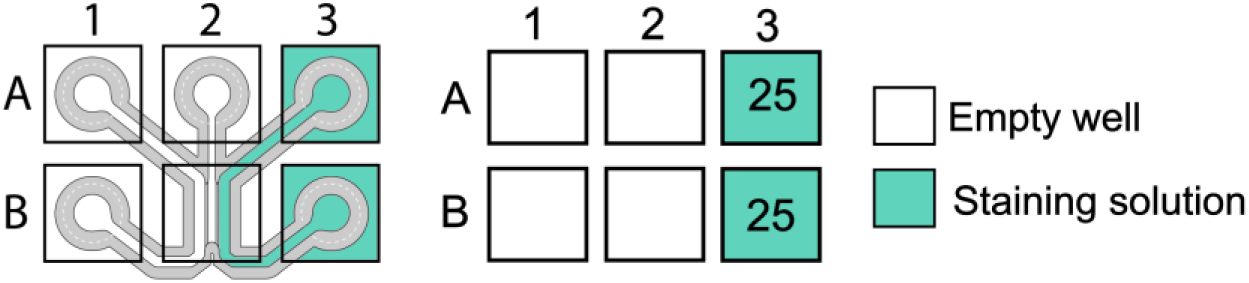
Pipetting scheme for ActinGreen and NucBlue staining.

**Figure 5.**
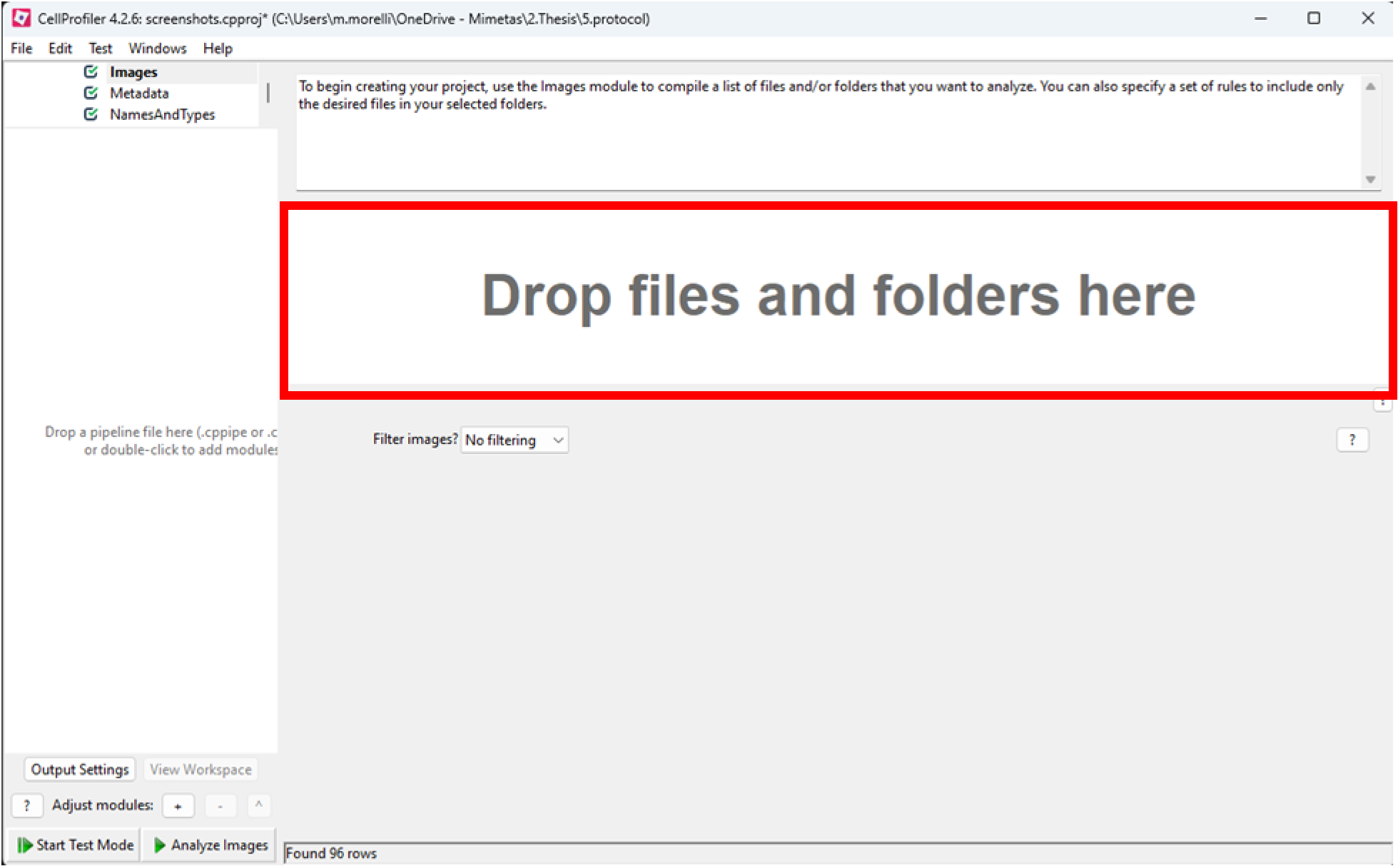
CellProfiler image loading interface. Go to image module and right click on the box highlighted in red and load images to be analyzed (drag and drop is also possible).

##### Cropping images

Before proceeding with the analysis, images need to be cropped to only select the channel in which there are cells:

1. click on “add a module” at the bottom left corner
2. select the “Crop” module in the “Image Processing category” and click “done”:
3. Once the crop module is added, it will appear as the first step of the workspace. Select it to open the details and adjust the following settings:
  a. Select the input image with the Name given in step 9
  b. Name the output image (ex: cropped_hoechst)
  c. Set cropping shape to rectange and add the following positions
    i. Left and right rectangle positions: 1350, end, absolute
    ii. Top and bottom rectangle positions: 0, end, absolute **Note**: The settings provided are for 3-lane 64 OrganoPlate format. For 3-lane-40 format, the cropping settings are the following:
      - Left and right rectangle positions: 0, end, absolute
      - Top and bottom rectangle positions: 150, 550, absolute
4. To test that the cropping settings are accurate, start the test mode, ensure the eye symbol is ticked and click on play.
5. A new window will pop up with the full image on the left and the cropped image on the right. The cropped image can now be referred to as “cropped_hoechst” in the pipeline, as specified in the crop settings.

**Note:** images can be rotated using the “FlipAndRotate” module in the “ImageProcessing” category.

**Note:** Cropped (and rotated) images can be saved using the “SaveImages” module in the “File Processing” category.

**Tip:** use the “Zoom” and the “Measure length” tools to measure the diameter of the nuclei. This will be helpful for the next step.

#### Hoechst positive objects identification

Hoechst is a dye commonly used to stain nuclei for cell counting purposes. In the quantification process, the goal is to establish a new object category named ‘hoechst_nuclei.’ All accepted objects per image will be classified under this category and can collectively be referred to by the ‘hoechst_nuclei’ name in subsequent analysis steps.

**Goal:** quantify Hoechst-positive nuclei

**Required:** Images of cells stained with Hoechst

To quantify Hoechst-positive objects:

1. Load and pre-process images as shown in section **Image pre-processing**.
2. add the “IdentifyPrimaryObjects” module from the “Object Processing” category and set the following settings
  a. Click yes on “Use advanced settings” and select the input image, which was generated in step (in our case the image is called cropped_hoechst)
  b. Name the object to be created (ex: hoechst_nuclei)
  c. Select the diameter range of the object. To determine the diameter range, activate **test mode**, ensure the display for the IdentifyPrimaryObjects module is activated by clicking the **eye icon** next to the module name, and click the **play icon** (as shown in **Figure 11**). In the display window, zoom in on a nucleus using the **Magnifier tool** and select the **Measure Length tool** from the toolbar. Click and drag over a nucleus to measure its diameter in pixels. Use these measurements to define the minimum and maximum diameter range for nuclei in the module settings.
  d. Ensure that objects outside the diameter range and object touching the border of the image are discarded
  e. Set “Method to distinguish clumped objects” to “Shape”
  f. Set “Method to draw dividing lines between clumped objects” to “Shape”
  g. Set “Fill holes in identified objects?” to “After both thresholding and declumping”
3. Use the test mode to check the output
  a. Make sure the “IdentifyPrimaryObjects” module is selected
  b. Start the test mode, and use the display mode to open a new window and access the following information:
    i. **Input image**: the raw image used
    ii. **Hoechst_nuclei outlines**: the object outlines with the accepted objects in green, the objects discarded because they touche the border in orange, and the objects discarded because they are outside the diameter range in purple.
    iii. **Hoechst_nuclei:** accepted objects
    iv. **Table with calculations** such as the number of accepted objects, their median diameter, the area covered by the objects, etc.
4. Hoechst identification is now done. Continue with **DRAQ7 object identification**, **Actin objects identification and quantification**, or see **Note:** While this protocol focuses on measuring area using the MeasureImageAreaOccupied module, additional modules such as MeasureObjectSizeShape, MeasureObjectIntensity, and MeasureImageIntensity can be used for deeper analysis. These modules can provide detailed insights into the geometry, intensity distribution within objects, or overall image intensity. If analyzing intensity, it is essential to work with sum projections of the images to ensure accurate intensity measurements. These modules and their analyses are beyond the scope of this protocol and will not be described here.
5. Exporting data if you are done.

**Figure 6.**
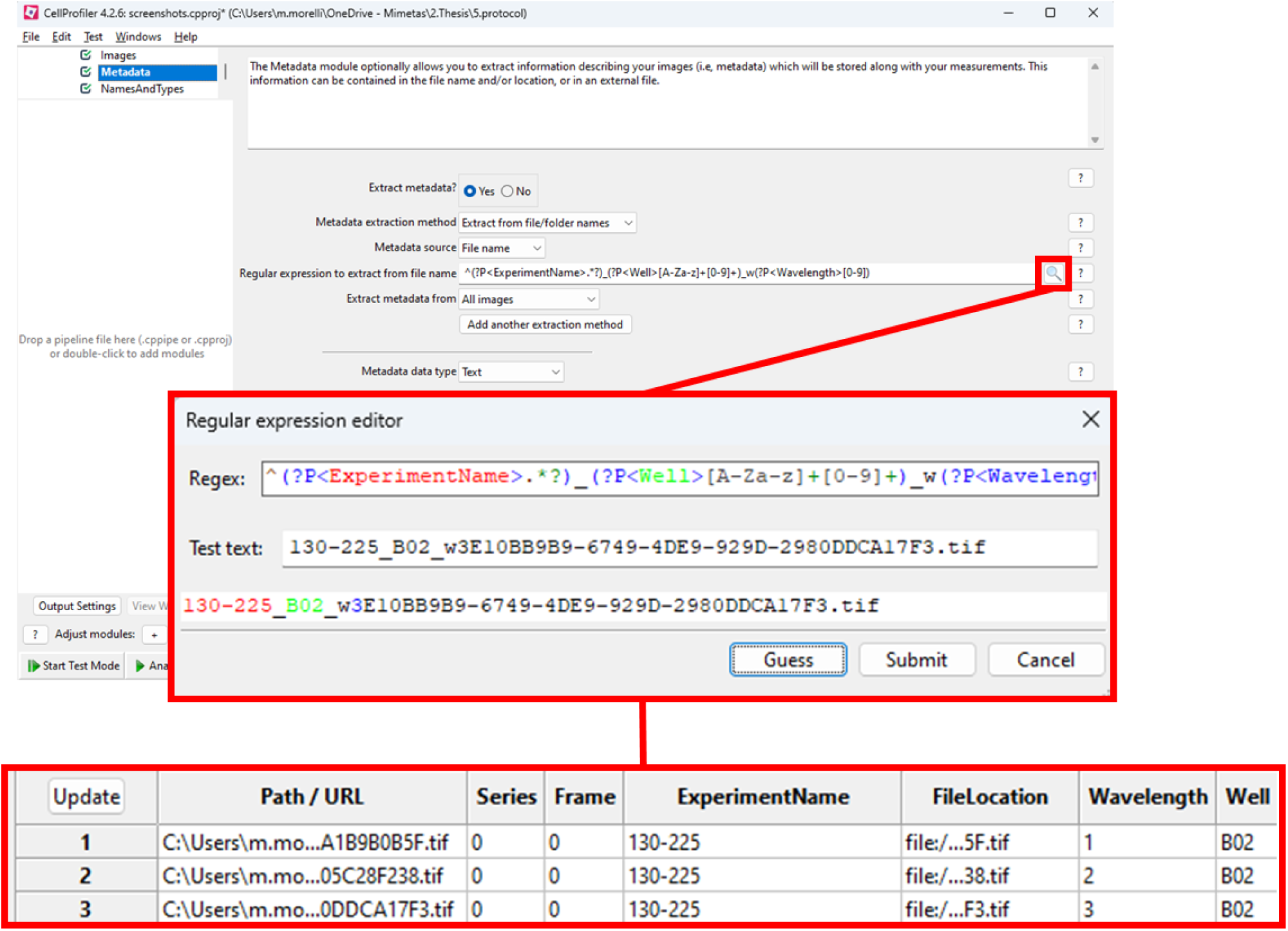
Extracting information from the image filenames using the Metadata module.

**Figure 7.**
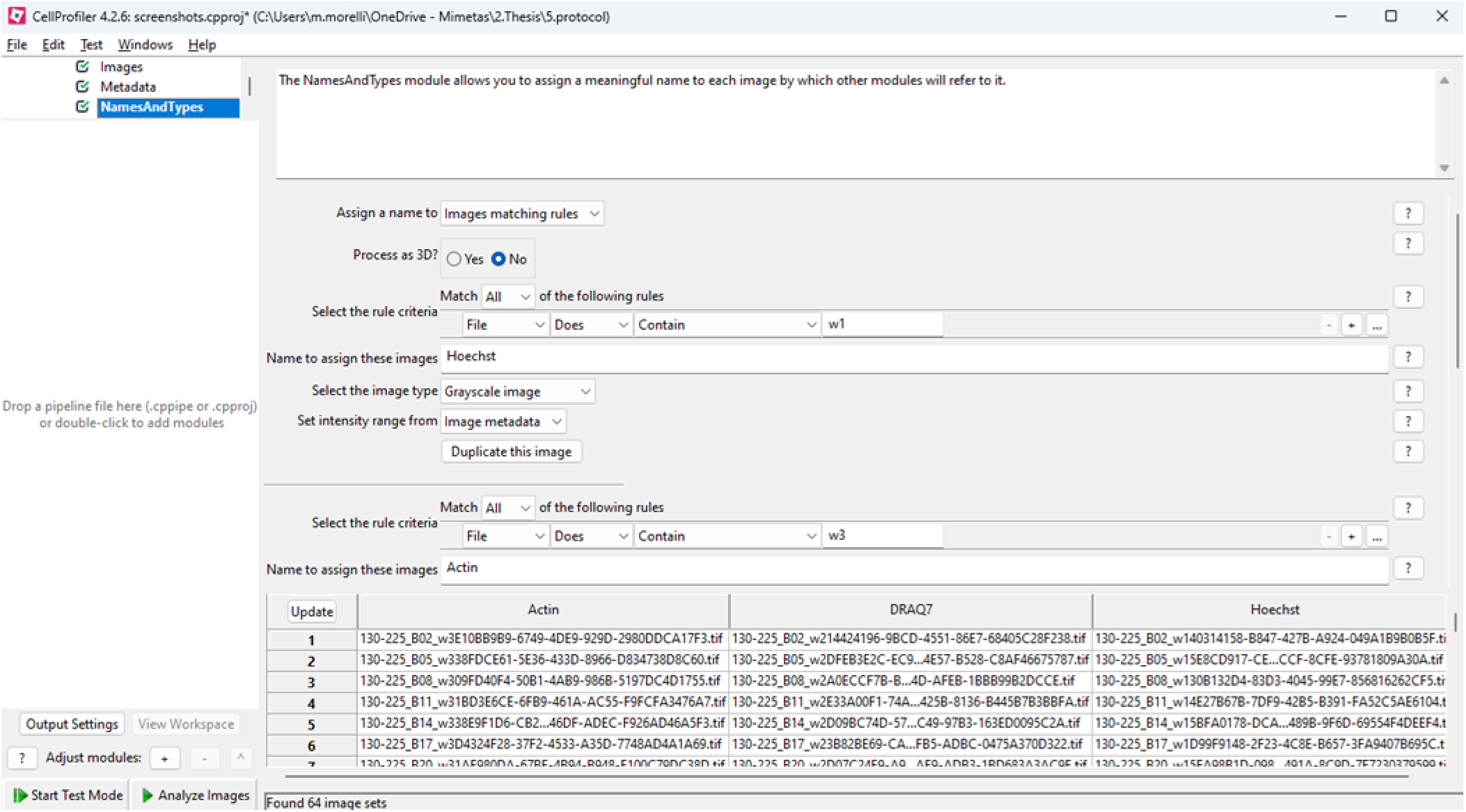
Assigning wavelength names to images based on filename using the NamesAndTypes module.

**Figure 8.**
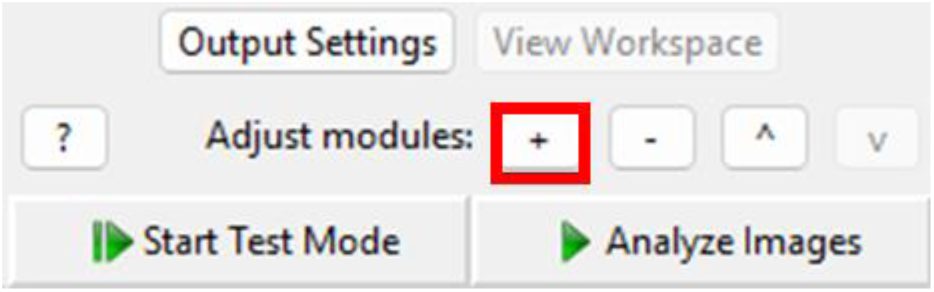
Add module button.

**Figure 9.**
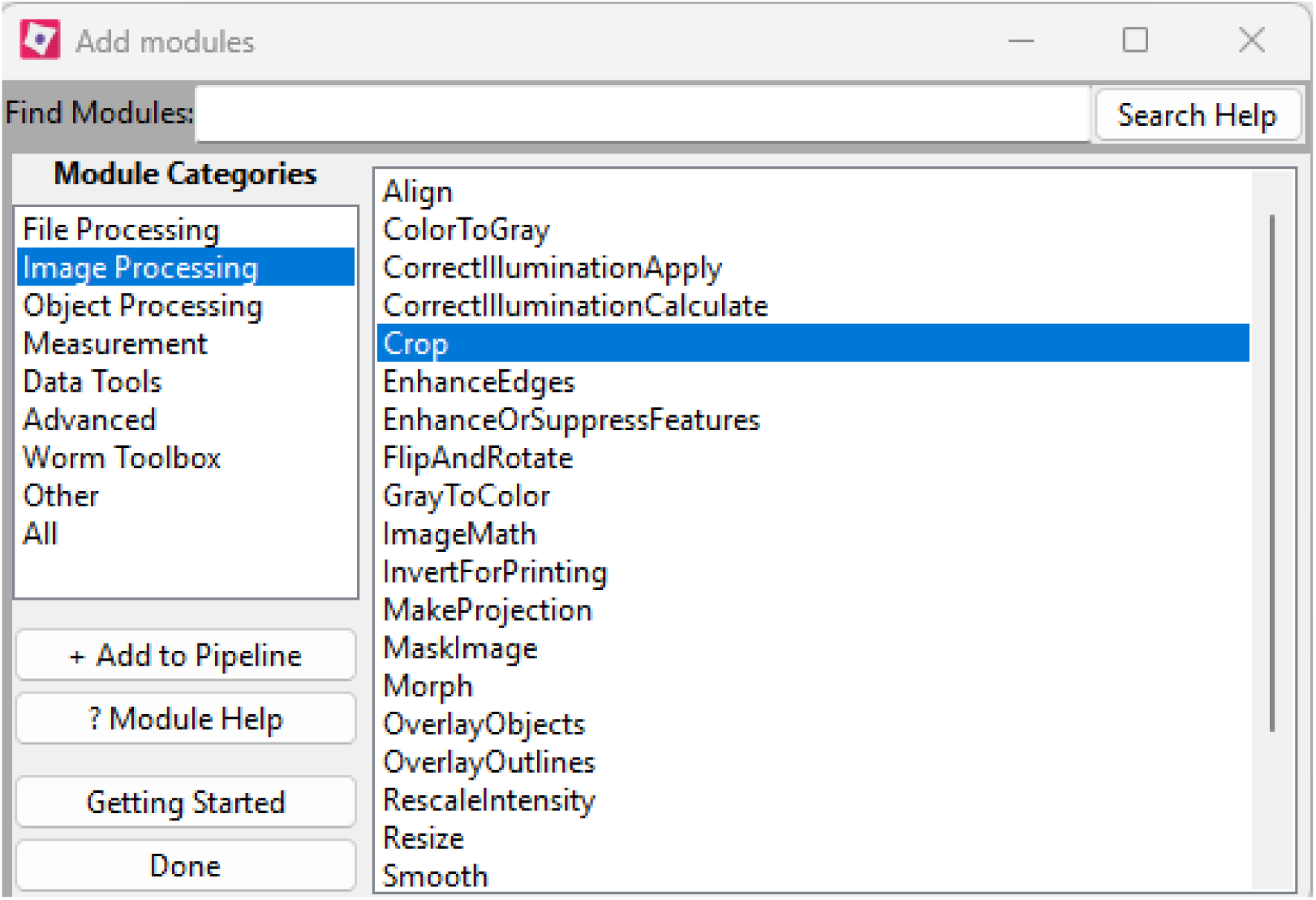
Add module window with the “Crop” module selected from the “Image Processing” category.

**Figure 10.**
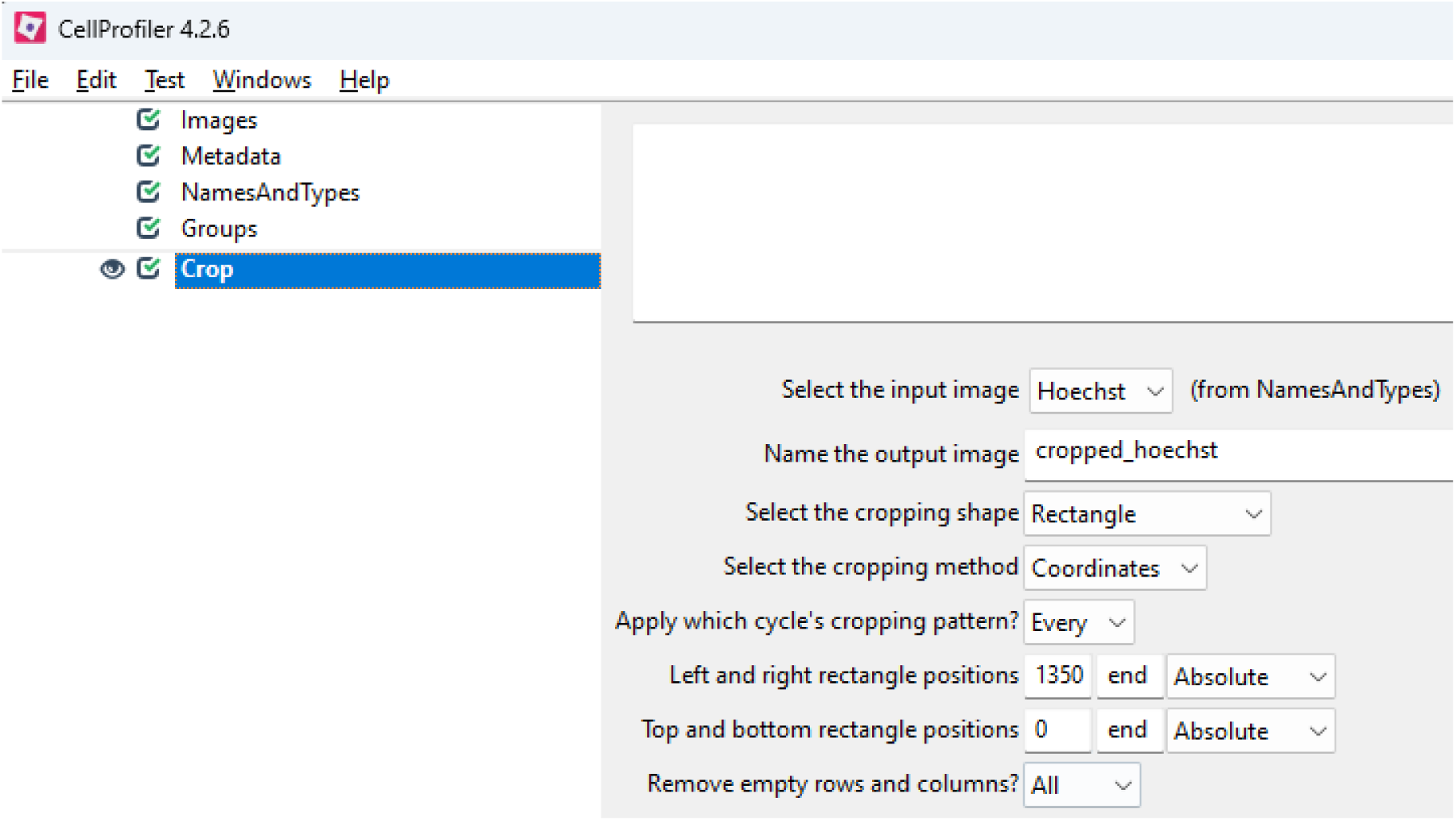
Crop module settings for 3-Lane-64 OrganoPlate with tubule in the right channel.

**Figure 11.**
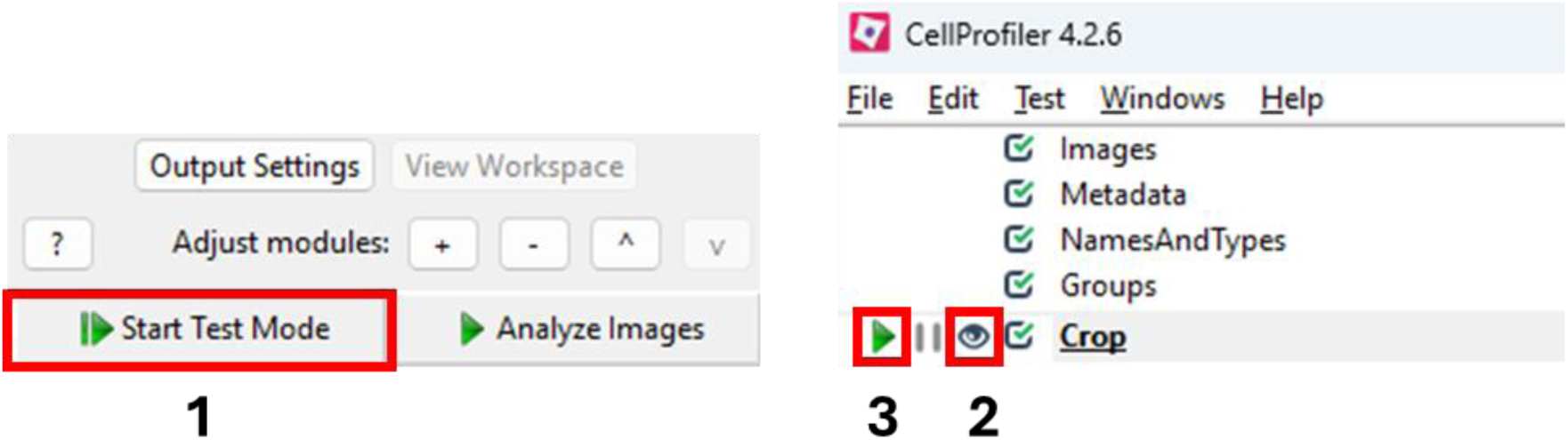
Sequence of steps to start the test mode and show the display of the selected module (crop module in this case).

**Note:** Outlines and masks of accepted objects can be saved for each image (see **Saving images, outlines and masks**).

**Note:** CellProfiler doesn’t save excluded objects and or subplots. To save a subplot (e.g. the bottom-left image on **Figure 15**) right-click and save it manually for every image.

**Figure 12.**
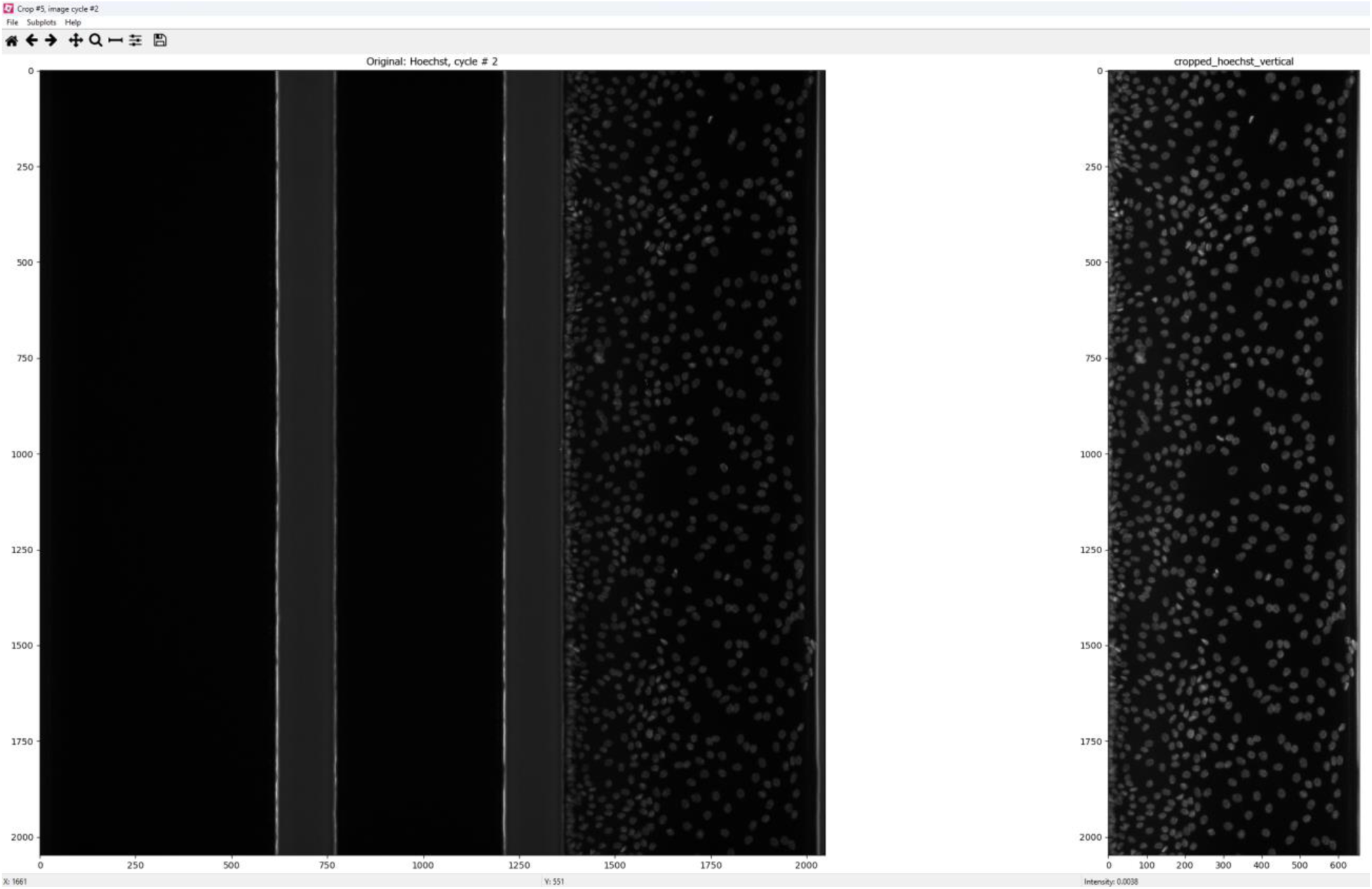
Crop module display window. Left is the original image and right is the cropped image. The cropped image dimensions are 425×1350 µM.

**Figure 13.**
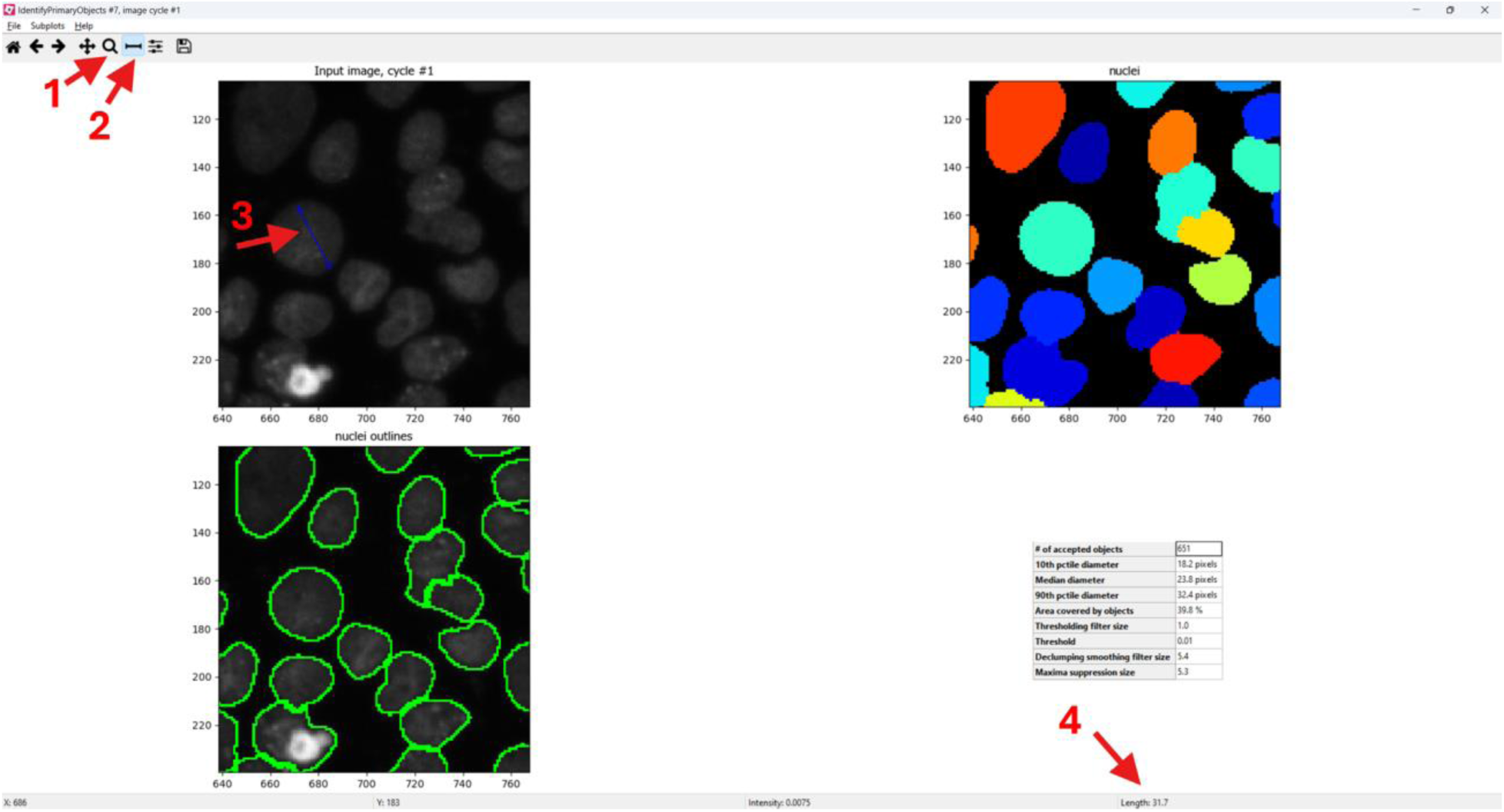
Measuring the diameter of a nuclei in the display window of the module. Zoom on the picture using the zoom tool (1), then select the measure length tool (2). Click and drag over a nucleus to measure its diameter in pixels (3) which will be shown at the bottom of right of the screen (4).

**Figure 14.**
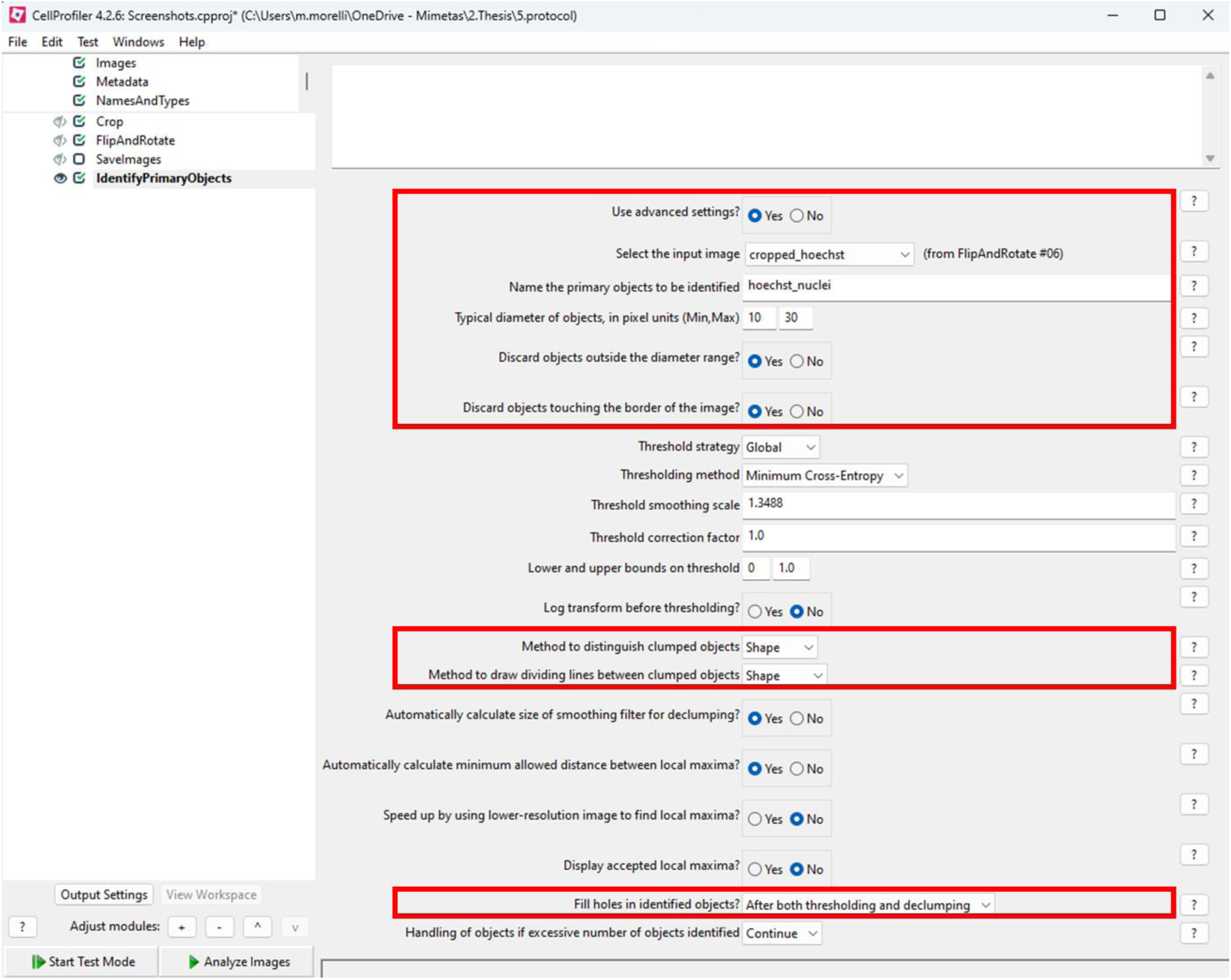
Settings of the IdentifyPrimaryObjects module for Hoechst-positive objects detection.

**Figure 15.**
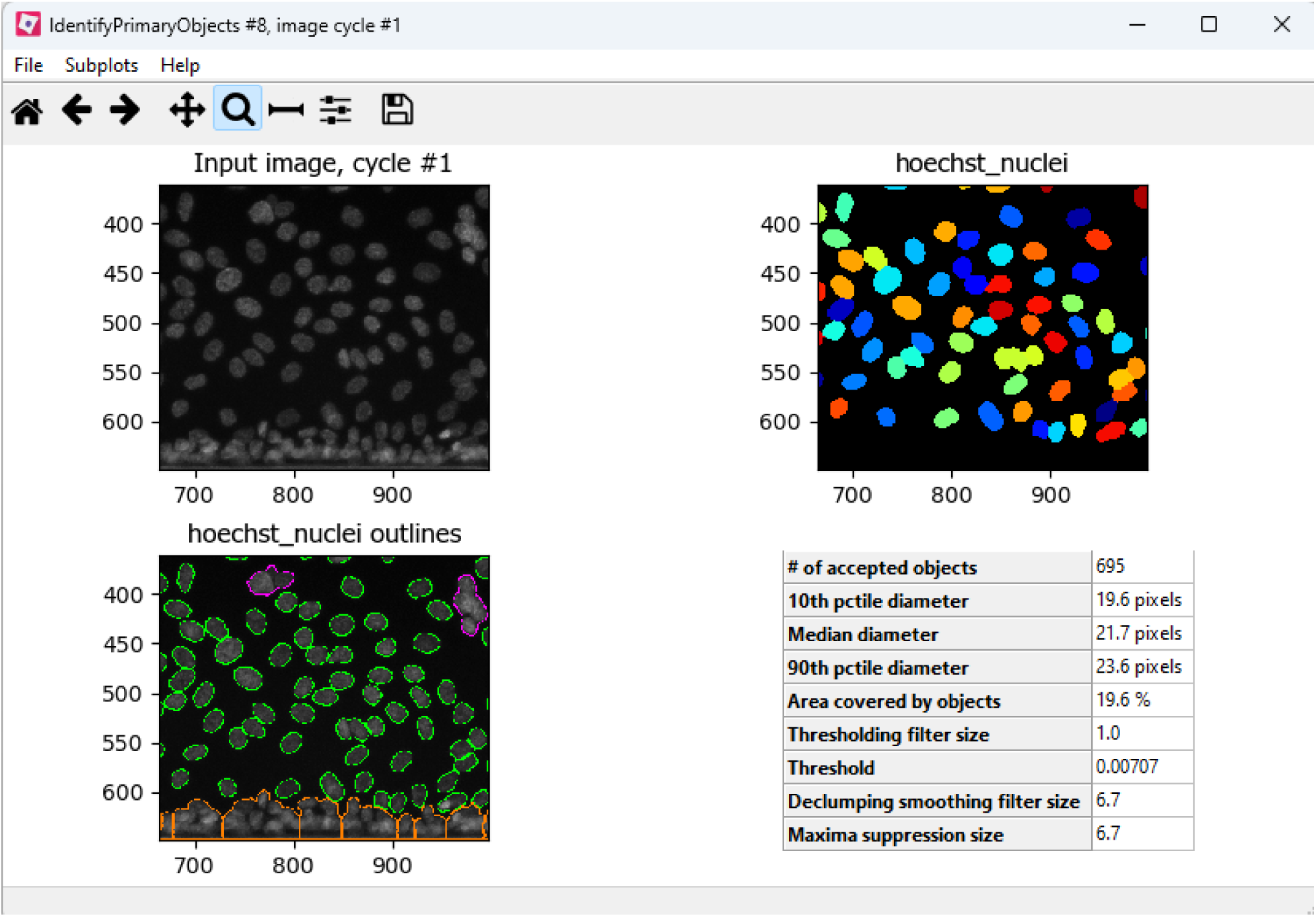
Hoechst-positive object identification. Accepted objects are shown in green, while excluded objects are shown in orange (if touching side of the image) or purple (if outside the diameter range). Note that the images have been zoomed in to better see the nuclei, but the exclusions and calculations apply to the whole image.

#### DRAQ7 object identification

DRAQ7 is a dye that selectively stains the nuclei of dead or permeabilized cells. In the quantification process, the goal is to establish a new object category named ‘draq7_nuclei.’ All accepted objects per image will be classified under this category and can collectively be referred to by the ‘draq7_nuclei’ name in subsequent analysis steps. Unlike Hoechst, which stains the nuclei of all cells regardless of their viability, DRAQ7 staining varies depending on the cell’s membrane integrity. Therefore, we recommend optimizing object quantification by using images of positive controls where DRAQ7 positive nuclei are present, and negative controls where DRAQ7 positive nuclei are rare or absent.

**Goal:** identify Draq7 positive nuclei and count them

**Required:** images from cells stained for Hoechst and Draq7+. Treated samples and corresponding negative control are recommended.

1. Load Hoechst and DRAQ7 images and do the necessary pre-processing, as shown in section **Image pre-processing**.
2. Identify Hoechst-positive objects, as shown in section **Hoechst positive objects identification**.
3. Add the “IdentifyPrimaryObjects” module from the “Object Processing” category
  a. Click yes on “Use advanced settings” and select the input image (cropped_draq7)
  b. Name the object to be created (ex: draq7_nuclei)
  c. Select the diameter range of the draq7 object
  d. Use the pixel intensity tool in test mode to set the lower bound of threshold (see **Figure 16** and **Figure 17**)
  e. Use the display mode (within test mode) to check the output.
  f. Save the outlines and masks

**Note:** Some images may not contain any DRAQ7-positive objects, and the quantification might lead to false-positives (see **Problem 2: false positives during object** identification).

**Figure 16.**
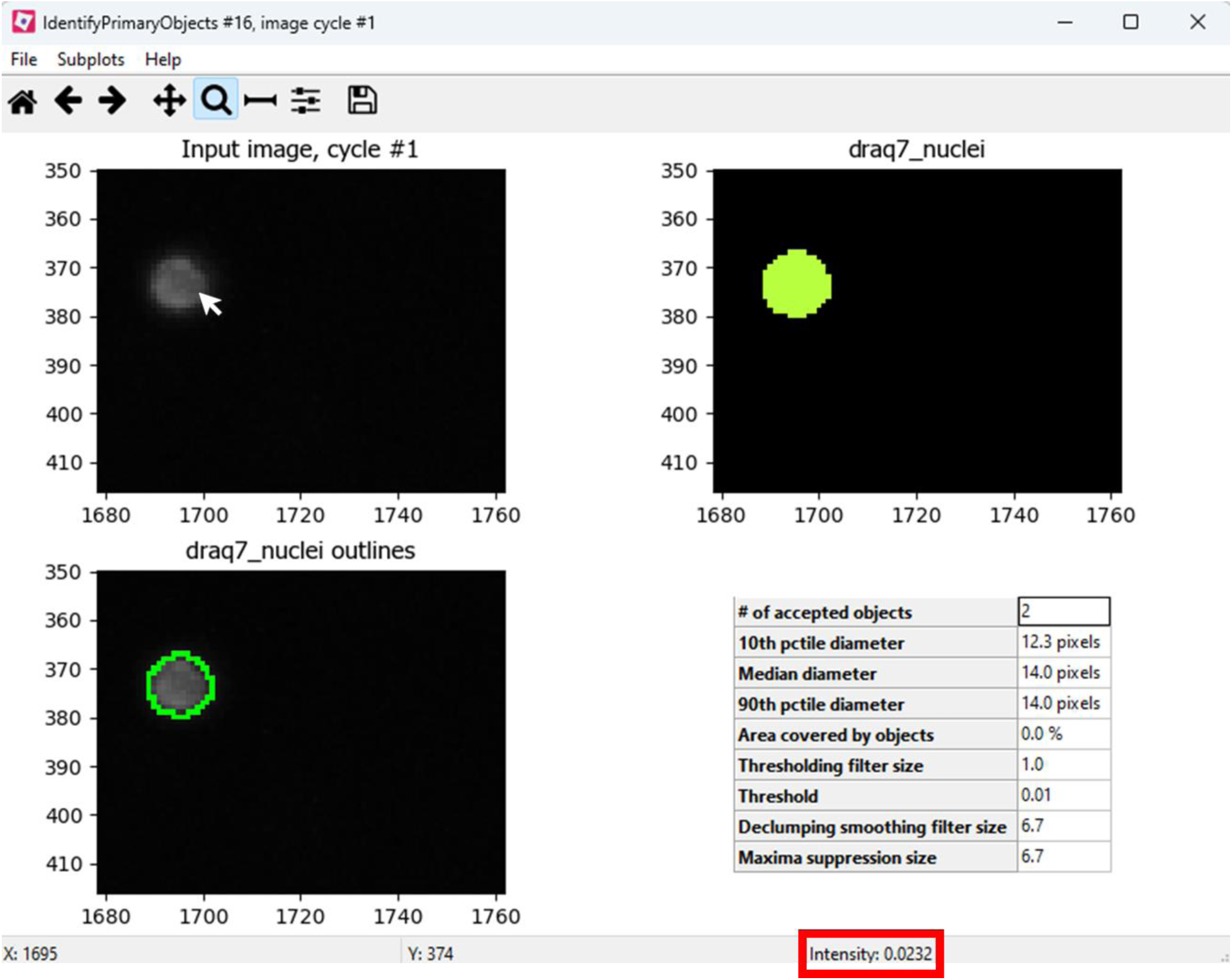
Pixel intensity tool showing the pixel intensity of an object.

**Figure 17.**
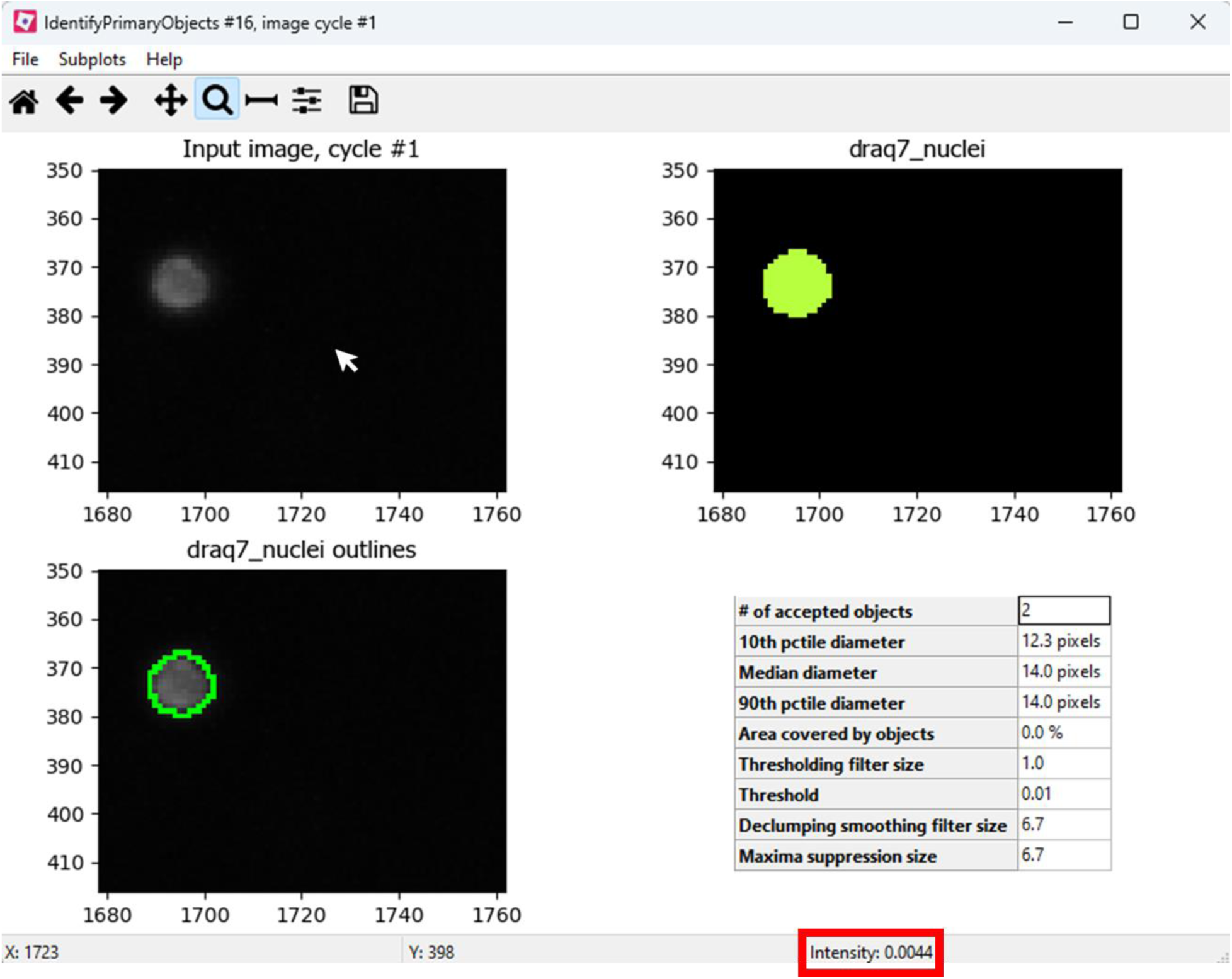
Pixel intensity tool showing the pixel intensity of the background.

### Calculating the percentage of DRAQ7-positive objects

To accurately quantify the percentage of DRAQ7-positive nuclei and minimize false positives, we will identify nuclei that are both DRAQ7 and Hoechst positive. This involves linking Hoechst-positive nuclei with corresponding DRAQ7-positive nuclei and selecting only those that are double-positive. These will then be categorized into a new object group named “Hoechst_draq7_nuclei”. To quantify Hoechst and DRAQ7 double positives:

1. Male sure you have identified Hoechst-positive and DRAQ7-positive objects (see **Hoechst positive objects identification and DRAQ7 object identification** sections)
2. Use the “RelateObjects” module in the “Object Processing” category.
  a. Set the “Parent objects” to “Hoechst_nuclei”
  b. Set the “Child objects” to “draq7_nuclei”
  c. Save the children with parents as new objects (objects with no parents will be discarded)
  d. Name the output objects (ex: Hoechst_draq7_objects)
3. Use the “ConvertObjectsToImage” and “SaveImages” modules to save the images (see Saving images, outlines and masks section)

To calculate the percentage DRAQ7-positive objects, the following operation will be performed:

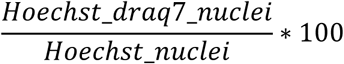

Hoechst_draq7_nuclei and Hoechst_nuclei count can be exported in a spreadsheet (see Exporting data into spreadsheets section) to do the calculation manually. Alternatively, the calculation can be done directly in the pipeline to create a new measurement that can be exported in a spreadsheet or displayed on images. To add the operation to the pipeline:

4. Use the “CalculateMath” module from the “Data Tools” category:
5. Name the output measurement (ex: percent_draq7)
  a. Set Operation to “Divide”
  b. Set the numerator to Image-> Count-> Hoechst_draq7_nuclei
  c. Set the denominator to Image-> Count-> Hoechst_nuclei
  d. Multiple the result by 100
  e. The percent_draq7 measurement can now be used to be displayed on images (using the DisplayDataOnImage module) or saved in the export spreadsheet.
6. DRAQ7 quantification is now done. Continue with **Actin objects identification and quantification**, or see **Exporting data** if you are done.

### Actin objects identification and quantification

Actin is a highly dynamic and structurally variable component of the cytoskeleton, making it challenging to quantify compared to nuclear markers like Hoechst and DRAQ7, which stain nuclei that are relatively uniform in shape and easier to identify. Actin’s shape and organization can vary significantly depending on cell type and treatment conditions, necessitating a tailored approach for its analysis. Here, we employ a straightforward method to quantify actin structures and measure their characteristics, such as area, perimeter, and the number of objects. These metrics provide valuable insights into cytoskeletal remodeling induced by toxins or other treatments.

Given the complexity of actin structures, quantification efforts are most effective when clear effects are observed in the images. To ensure resources are focused on biologically relevant changes, it may be practical to first visually inspect the images and prioritize quantification for conditions showing discernible differences from controls. This approach streamlines analysis while preserving the reliability of results.

**Goal:** identify actin positive objects and measure their properties such as number, area, perimeter.

**Required:** images from cells stained for Hoechst and Actin, including samples with treatment and a corresponding negative control for comparison.

1. Load Hoechst and ACTIN images and do the necessary pre-processing, as shown in section **Image pre-processing**
2. Identify Hoechst-positive objects, as shown in section **Hoechst positive objects identification**.
3. Use the IdentifyPrimaryObjects module:
  a. Click yes on “Use advanced settings” and select the input image (cropped_actin)
  b. Name the object to be created (ex: actin)
  c. Select the diameter range of the actin objects. Use the measure length tool if needed (**Figure 13**).
  d. Ensure “Discard objects outside of diameter range?” is set to **Yes**
  e. Ensure “Discard objects touching the border of the image?” is set to **Yes**. **Note:** Set this option to **Yes** to avoid false-positive objects caused by autofluorescent microfluidic structures. If actin structures near the border are critical, this can be set to **No**, but be aware of potential inaccuracies from partial objects.
  f. Set the lower bound of threshold: in test mode, activate the display in the IdentifyPrimaryObjects module and click the play icon to view the results. Hover over the input image with your mouse to check pixel intensity values displayed at the bottom of the window (see **Figure 16** and **Figure 17**). Use these values to determine the lower bound of the threshold. Adjust the threshold as needed and click play again to see the updated results. **Note:** Unlike nuclei quantification, where the lower bound of threshold is typically set just above the background, defining the lower threshold for actin can be more challenging and subjective. This is because actin staining is ubiquitous, and the goal here is to identify brighter clumps of actin rather than the general background signal. Aim to set the threshold to exclude the normal actin signal while capturing distinct, higher-intensity regions indicative of clumps. Examples are given in **Figure 20** and **Figure 21**.
  g. Set “Method to distinguish clumped objects” to “None”. **Note:** This setting avoids unnecessary splitting of actin structures, which are highly variable and not tied to individual cells like nuclei.
4. Save the outlines and masks (see **Saving images, outlines and masks** section) and check them to ensure the identification is successful before moving to the next step.

**Figure 18.**
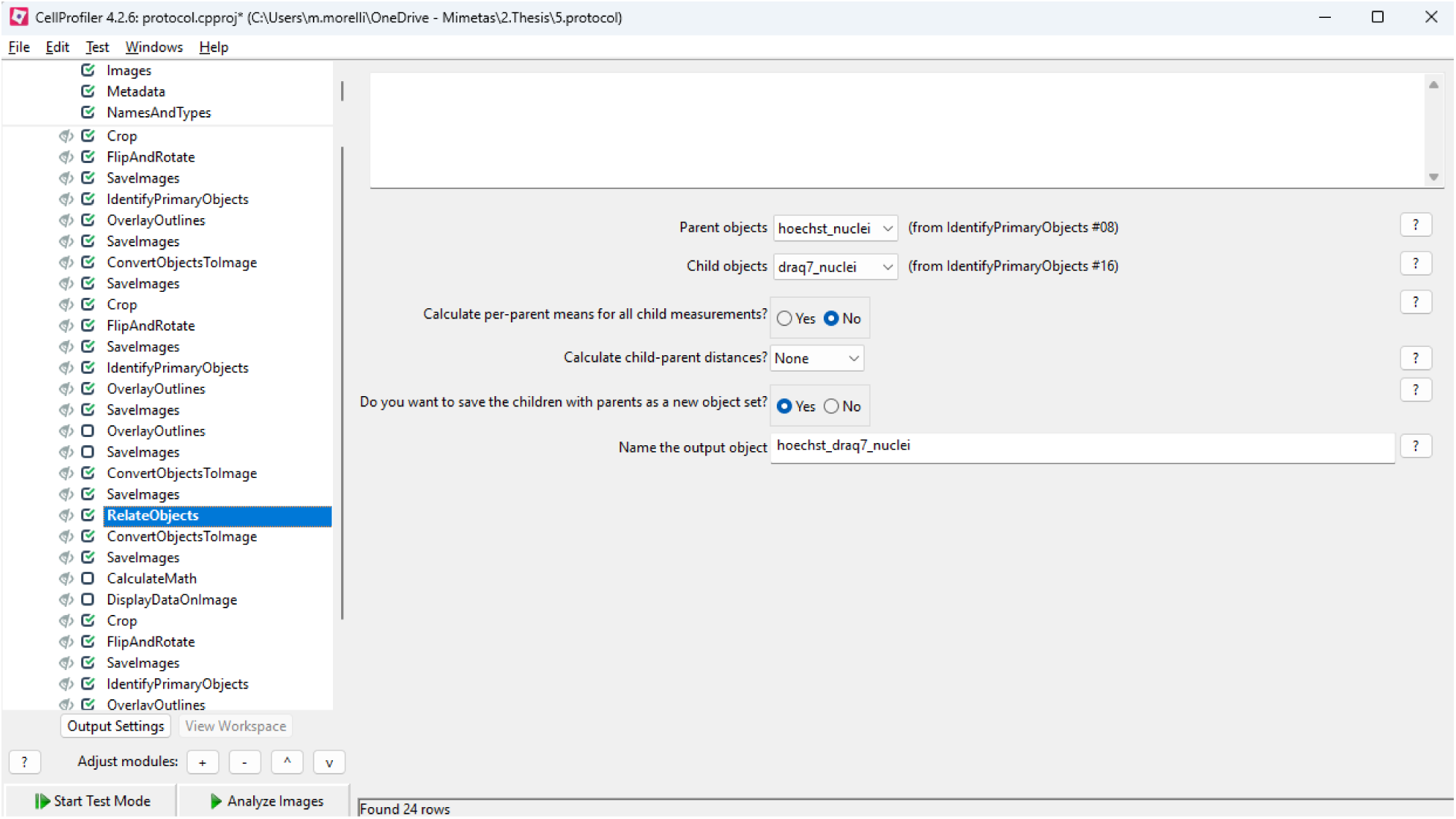
RelateObjects module settings for DRAQ7 quantification. Parent objects is set as “Hoechst_nuclei”, Child objects set as “draq7_nuclei”, and new output object named “hoechst_draq7_nuclei”.

**Figure 19.**
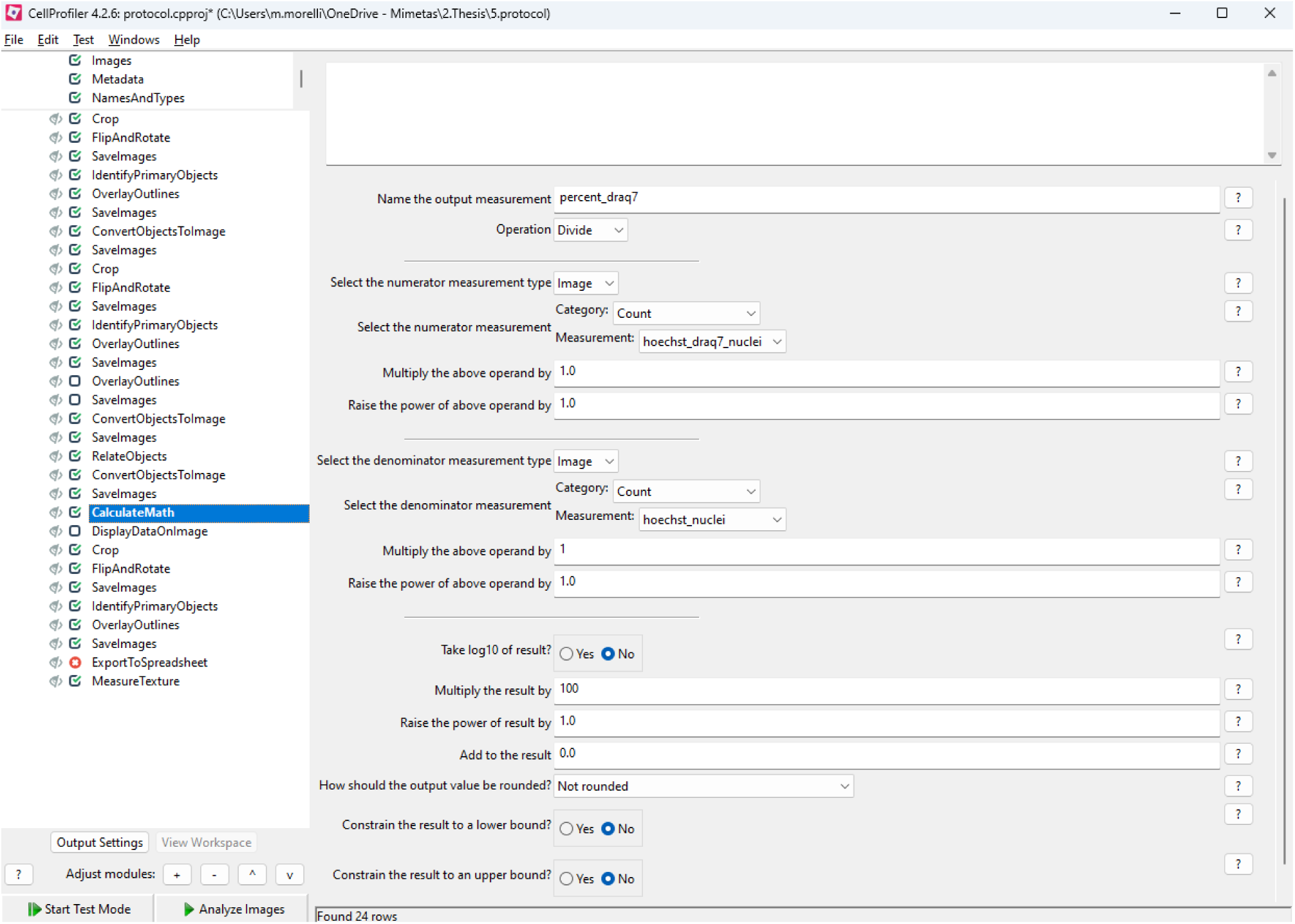
CalculateMath module set to calculate the percentage of draq7 positive objects. The. output measurement (percent_draq7) is obtained by dividing Hoechst_draq7_nuclei by Hoechst_nuclei and multiplying the result by 100.

**Figure 20.**
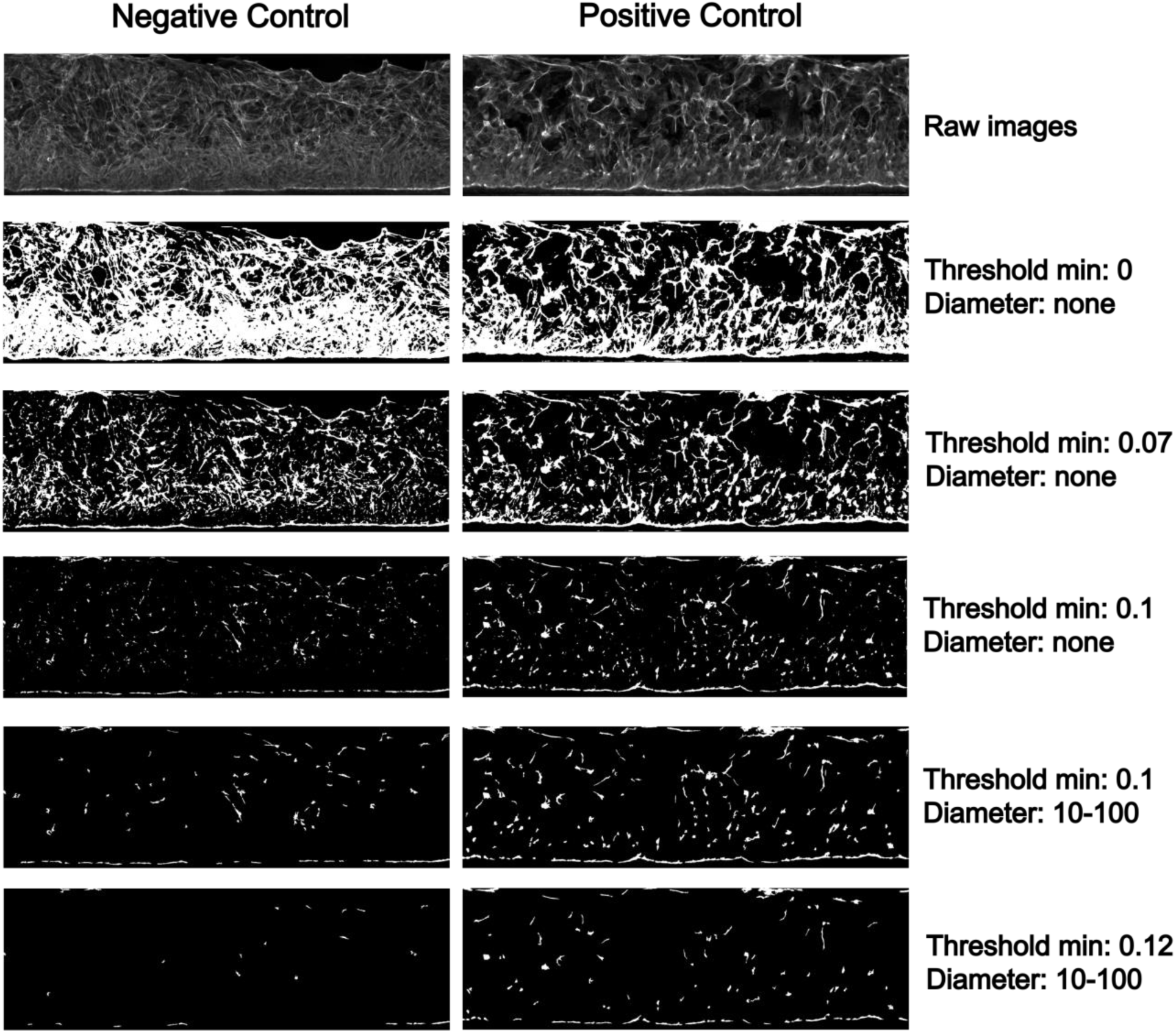
Identification of actin objects through variations in threshold and diameter parameters. Images are from OrganoReady® Colon Organoids and the positive control is exposed to candidalysin (75uM, 2h incubation). “Threshold min” is the lower bound of threshold, and “Diameter” is the diameter range, both derived from the IdentifyPrimaryObjects module. Images are 425×1350 µM.

**Figure 21.**
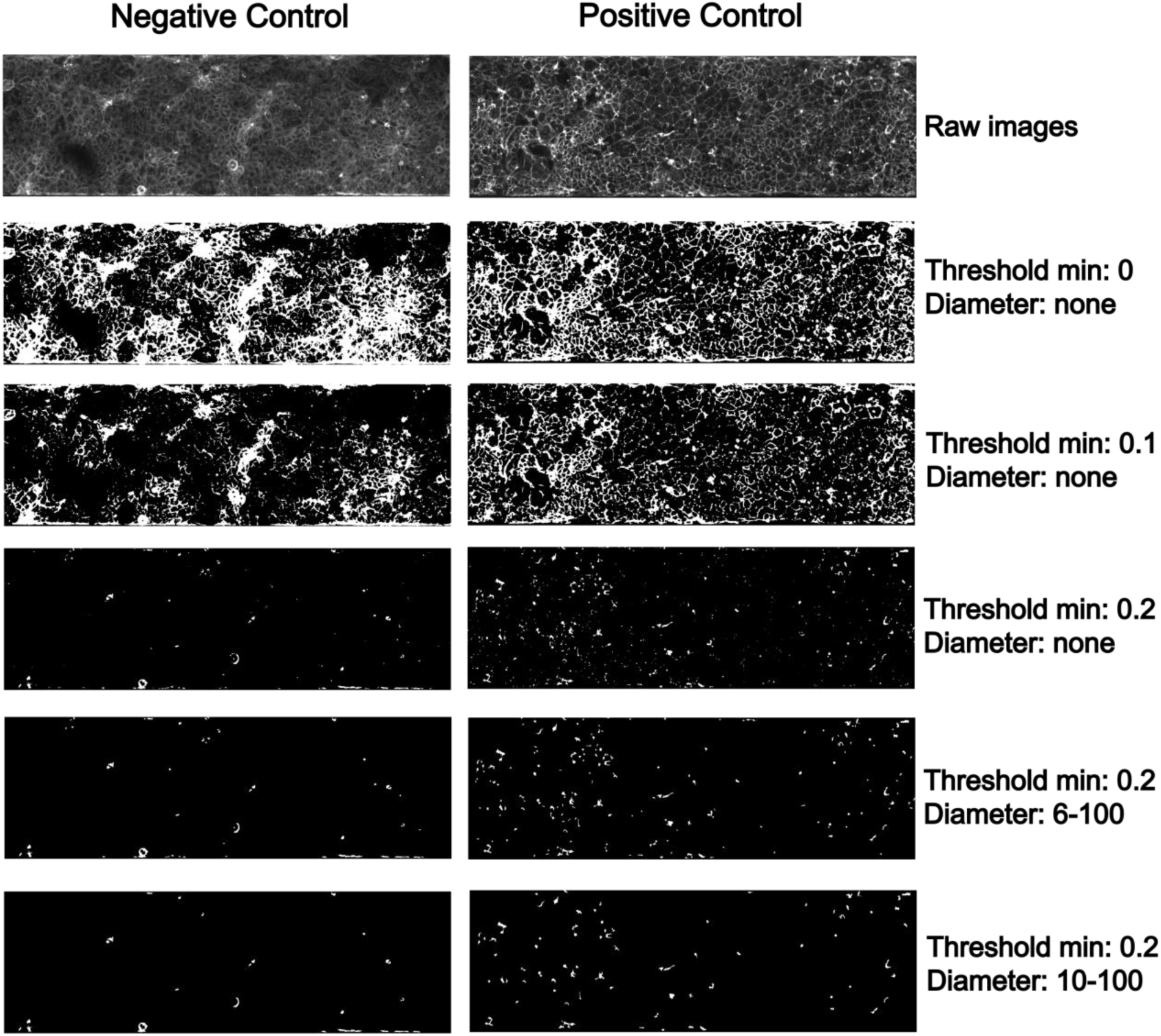
Identification of actin objects through variations in threshold and diameter parameters. Images are from OrganoReady® Colon Caco-2 and the positive control is exposed to nigericin (5 µg.ml, 5h incubation). “Threshold min” is the lower bound of threshold, and “Diameter” is the diameter range, both derived from the IdentifyPrimaryObjects module. Images are 425×1350 µM.

Once the actin objects are identified, the next step is to quantify the parameters of interest. In this analysis we will measure the area using the MeasureImageAreaOccupied module:

1. Open the MeasureImageAreaOccupied module:
  a. Select “Objects” in the drop-down list next to “Measure the area occupied by”
  b. Select the primary object of interest
2. **Optional:** use CalculateMath module to divide AreaOccupied by the number of actin objects to get the average object area
3. Actin quantification is now done. Continue with **DRAQ7 object identification**, or see **Exporting data** if you are done and want to export your data into spreadsheets or save images and masks.

**Note:** While this protocol focuses on measuring area using the MeasureImageAreaOccupied module, additional modules such as MeasureObjectSizeShape, MeasureObjectIntensity, and MeasureImageIntensity can be used for deeper analysis. These modules can provide detailed insights into the geometry, intensity distribution within objects, or overall image intensity. If analyzing intensity, it is essential to work with sum projections of the images to ensure accurate intensity measurements. These modules and their analyses are beyond the scope of this protocol and will not be described here.

## Exporting data

### Saving images, outlines and masks

Saving masks, outlines, and images in a CellProfiler pipeline is useful for quality control, troubleshooting, and documentation, as it allows for verification of segmentation accuracy and provides a visual reference for analysis results. We recommend saving them to check the quantification of each object before moving further with the analysis:

- To save outlines, use the “OverlayOutlines” module in the “Image Processing” category, then use the SaveImage module.
- To save the masks of the Hoechst_nuclei objects, use the “ConvertObjectToImage” module in the “ObjectProcessing” category, then use the SaveImage module.

Images can’t be saved in test mode, so the analysis needs to be run in regular mode to ensure all images and results are properly saved.

**Tip:** in the “Output File Location”, use the option “Default Output Folder sub-folder” to organize your data into sub-folders that you can name yourself.

#### 1.1.1.1 Exporting data into spreadsheets

1. Use the “ExportToSpreadsheet” module in “File Processing” category to generate csv file with the data.
2. Click **Yes** on “Select measurements to export”
3. Click on “Press button to select measurements”
4. Select the measurements you need and ensure all required parameters for analysis are included (as shown in **Figure 22**):
  a. Image Measurements
    i. **AreaOccupied** (generated by the MeasureImageAreaOccupied module <u>for</u> <u>the actin quantification</u>).
    ii. **Count** (automatically generated when adding the IdentifyPrimaryObject module, useful for actin and draq7 quantification).
  b. Metadata
    i. **ExperimentName** to track independent experiments (independent plates)
    ii. **Well** to track conditions (independent chips) **Note:** when using the CalculateMath module make sure to select the measurement for export (Select Measurements è Image è Math è your_measurement)
5. Select “Image” in “Data to export”

**Figure 22.**
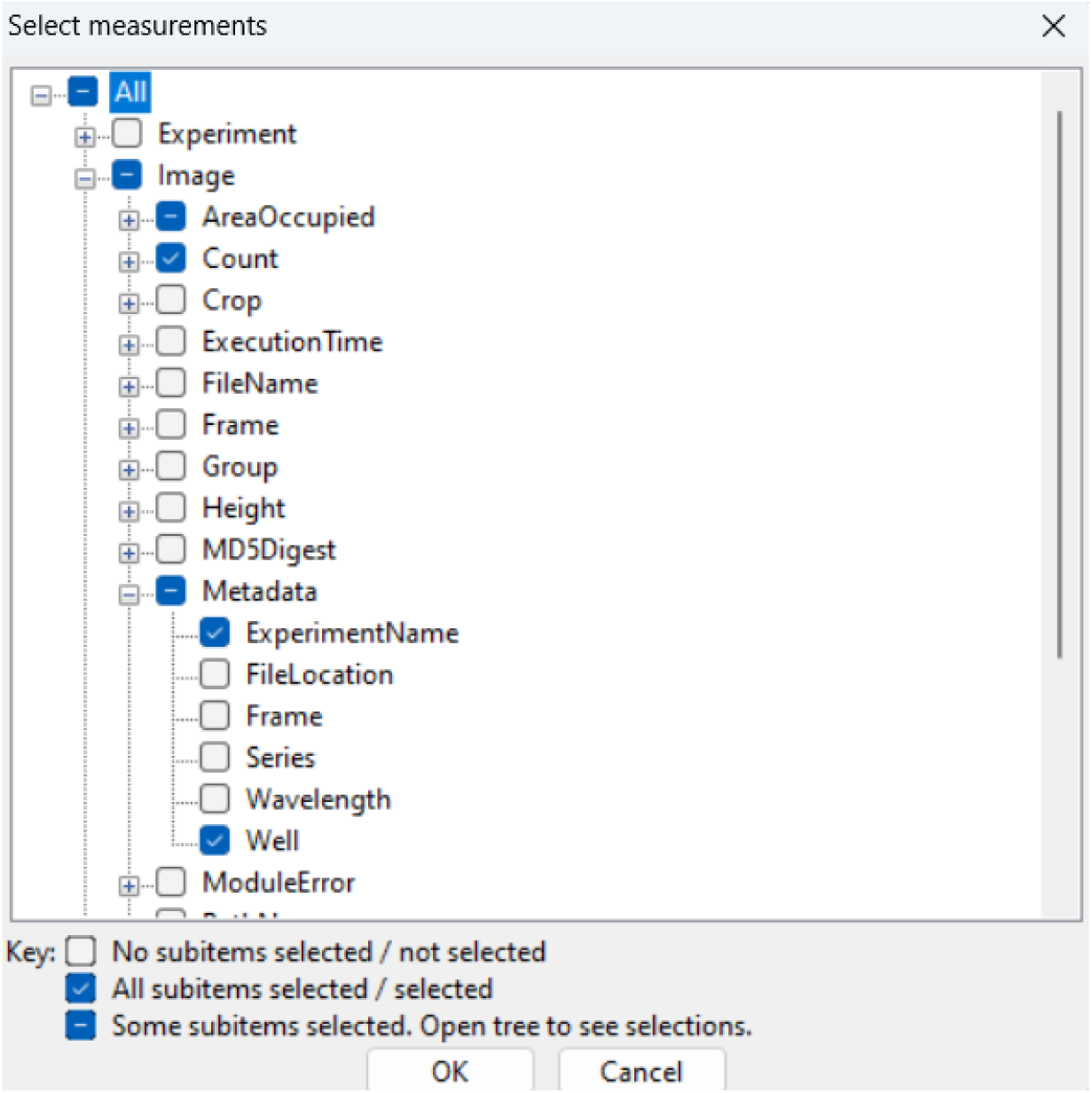
Pop up window to select measurements to export in spreadsheets.

**Note**: by default CellProfiler will generate multiple Excel sheets:

- **Experiment**: Contains general information, such as the software version and input/output folder details.
- **Image**: Provides a summary workbook with the average of relevant parameters per image (each row corresponds to an image).
- **Object-specific Sheets**: Generates one sheet per primary object, containing detailed measurements for each object (each row represents an object). These are particularly useful for deeper analyses using modules like MeasureObjectSizeShape, which calculate detailed parameters for individual objects.

In our case, the “Image” sheet, which averages selected parameters per image, is sufficient for analysis.

6. Set “Use the object name for the file name?” to no and name your dataset
7. Click on “Analyze Images” on the bottom left corner

**Tip:** make sure you disable the display window for all the modules before launching the analysis, otherwise the display window will pop up for each image set when analyzing the data.

### Data analysis

Output measurements can be found in the image excel sheet generated in the output folder after the images have been analyzed. An example output is given for actin quantification (**Table 2**). A dataset including the raw images, CellProfiler project files, and example outputs for ACTIN and DRAQ7 quantification, is provided as a supplemental dataset.

**Table 2.** Example output from actin quantification.

| AreaOccupied_Area<br>Occupied_actin | Count_actin | Count_nuclei | ImageNu<br>mber | Metadata_Expe<br>rimentName | Metadata_Well |
| --- | --- | --- | --- | --- | --- |
| 7076 | 31 | 619 | 1 | 130-226 | E02 |
| 7087 | 27 | 508 | 2 | 130-226 | E05 |
| 3609 | 19 | 481 | 3 | 130-226 | E08 |
| 3167 | 17 | 510 | 4 | 130-226 | E11 |
| 9917 | 27 | 584 | 5 | 130-226 | E14 |
| 3341 | 19 | 608 | 6 | 130-226 | E17 |
| 4113 | 23 | 518 | 7 | 130-226 | E20 |
| 3160 | 14 | 636 | 8 | 130-226 | E23 |
| 18063 | 48 | 537 | 9 | 130-226 | H02 |
| 18172 | 57 | 606 | 10 | 130-226 | H05 |
| 15639 | 58 | 528 | 11 | 130-226 | H08 |
| 14062 | 60 | 500 | 12 | 130-226 | H11 |
| 2723 | 16 | 516 | 13 | 130-226 | H14 |
| 9395 | 36 | 691 | 14 | 130-226 | H17 |
| 1588 | 10 | 660 | 15 | 130-226 | H20 |
| 1989 | 10 | 643 | 16 | 130-226 | H23 |
| 27625 | 78 | 606 | 17 | 130-226 | K02 |
| 18799 | 51 | 514 | 18 | 130-226 | K05 |
| 19857 | 54 | 569 | 19 | 130-226 | K08 |
| 11210 | 42 | 454 | 20 | 130-226 | K11 |
| 17549 | 42 | 654 | 21 | 130-226 | K14 |
| 9944 | 31 | 565 | 22 | 130-226 | K17 |
| 15645 | 69 | 679 | 23 | 130-226 | K20 |
| 3621 | 15 | 560 | 24 | 130-226 | K23 |
| 41585 | 86 | 655 | 25 | 130-226 | N02 |
| 34948 | 82 | 601 | 26 | 130-226 | N05 |
| 20828 | 81 | 540 | 27 | 130-226 | N08 |
| 40674 | 112 | 674 | 28 | 130-226 | N11 |
| 24621 | 60 | 615 | 29 | 130-226 | N14 |
| 4616 | 23 | 577 | 30 | 130-226 | N17 |
| 20667 | 58 | 691 | 31 | 130-226 | N20 |
| 11282 | 29 | 491 | 32 | 130-226 | N23 |

To account for differences in cell number across conditions, it is essential to normalize readouts by the number of nuclei per image. This ensures that observed changes reflect per-cell effects rather than variations in seeding density, treatment-induced cell loss, or imaging artifacts. In this protocol, we extract the total number of nuclei (Count_nuclei) during image analysis and perform post hoc normalization of all readouts based on this value.

Using the data shown in **Table 2**, normalized output measurements would be calculated as:

1. AreaOccupied_AreaOccupied_actin / Count_nuclei
1. Count_actin / Count_nuclei

**Table 3** references studies using this protocol and the controls used for each study.

**Table 3.**
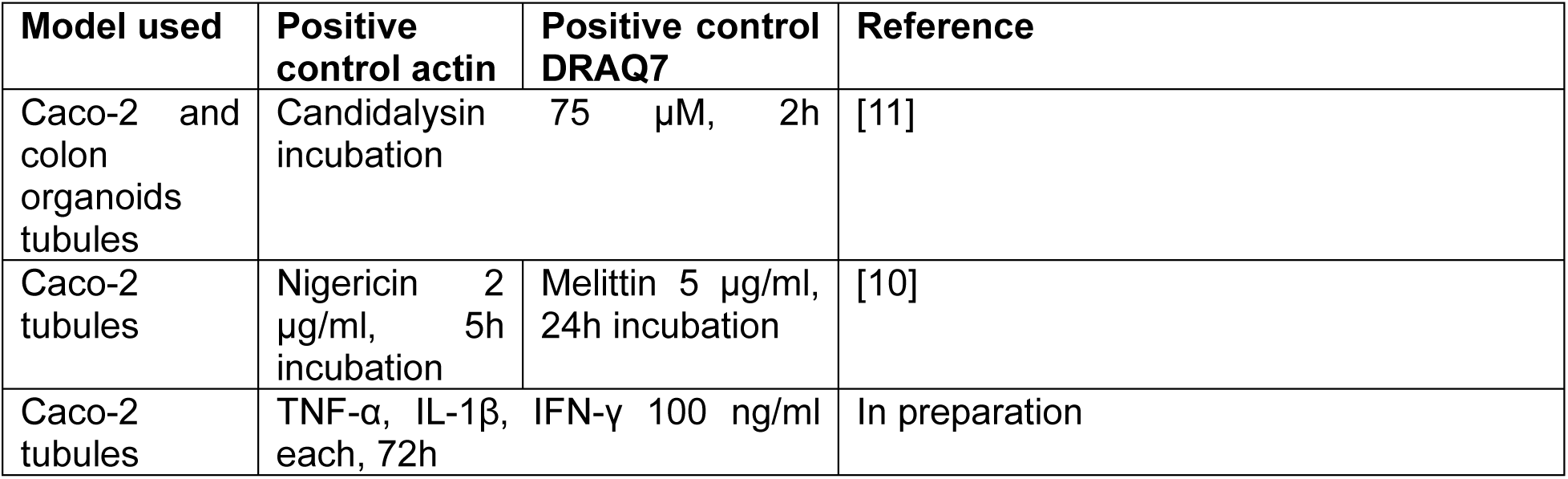
References using this protocol.

### Troubleshooting

#### Problem 1: tubule detachment

**Issue:** Tubule detachment after treatment disrupts the model’s structure and can interfere with accurate quantification.

**Cause:** Excessive cytotoxicity leading to cell death and loss of adhesion.

**Solution**:

- Reduce concentration: Perform dose-response analysis to determine the minimum effective concentration
- Shorten incubation: test reduced exposure times to limit cytotoxicity.

**Tip:** Monitor cell viability using light imaging and/or cytotoxicity assays to optimize conditions and prevent detachment.

### Problem 2: false positives during object identification

**Issue:** False positives may occur during analysis if no DRAQ7-positive objects are present (see **Figure 23**). This can happen in a negative control or a treatment with no effect on permeability.

**Figure 23.**
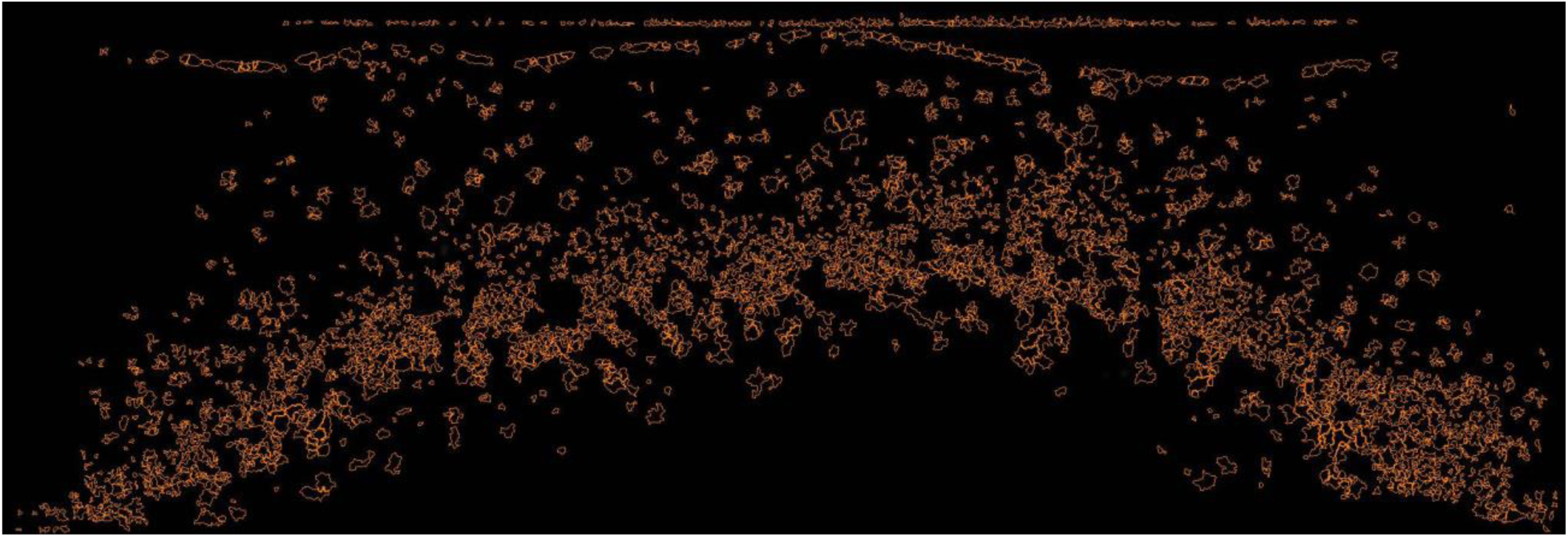
Example of false positive objects in an image with no signal. When the lower bound threshold is set to zero, a false positive can occur if the thresholding algorithm attempts to distinguish between foreground and background, but no objects are present in the image to define a foreground. This situation can lead to incorrect positive objects identifications, as shown with the orange outlines. Image is 425×1350 µM.

**Cause:** The thresholding algorithm incorrectly identifies background as signal

**Solution:** Increase the lower bound of threshold:

1. Use the pixel intensity tool in the image analysis software.
2. Hover the mouse over the image to display pixel intensities at the bottom left corner of the window.
3. Measure the pixel intensity of both the background and the signal.
4. Set the lower threshold to a value between these two measurements.

**Tip:** in the IdentifyPrimaryObject module, a setting called “Handling of objects if excessive number of objects identified” allows you to set a maximum number of objects and discard an image if the number of identified objects is higher. This setting is particularly useful when analyzing samples with consistent expected object numbers, as it allows you to focus on reliable data without manual filtering.

## Reagents

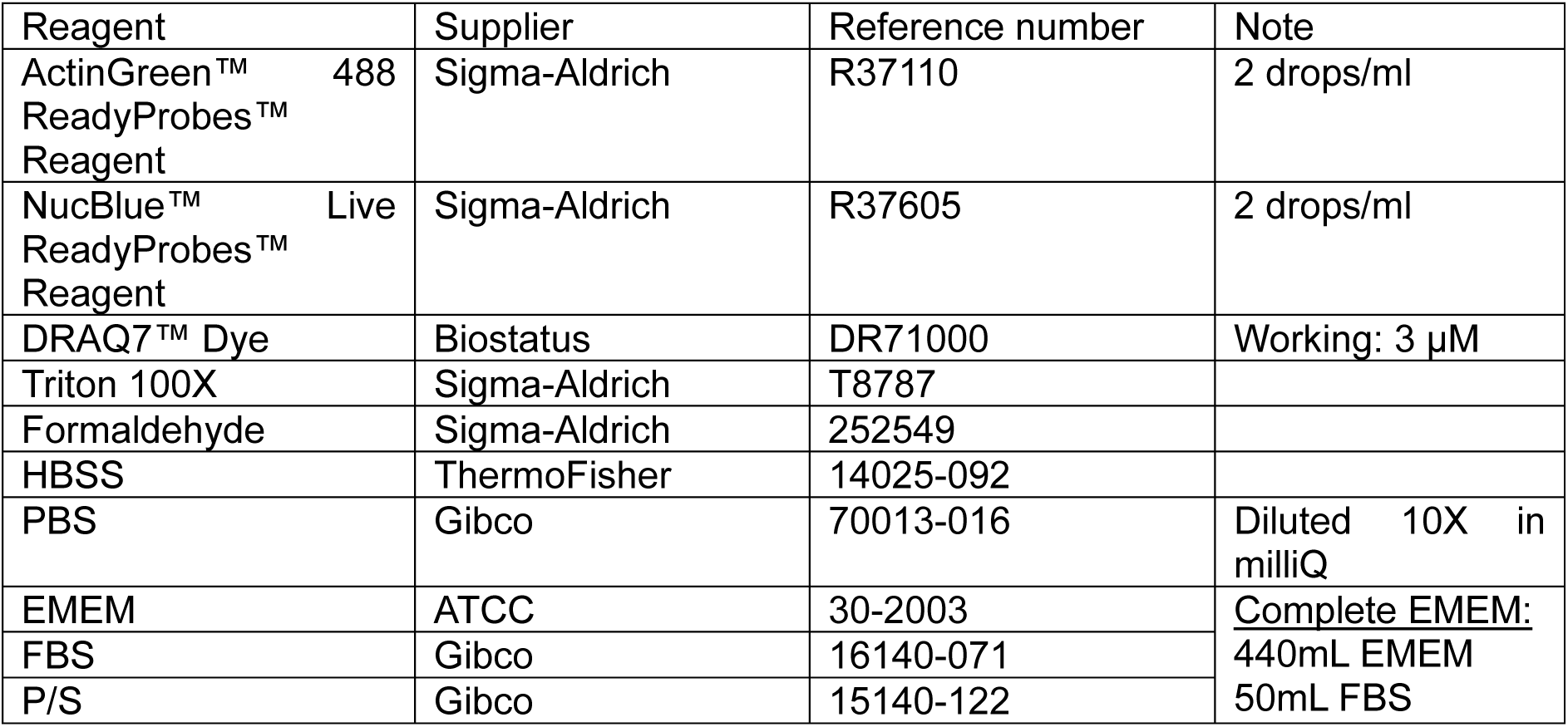
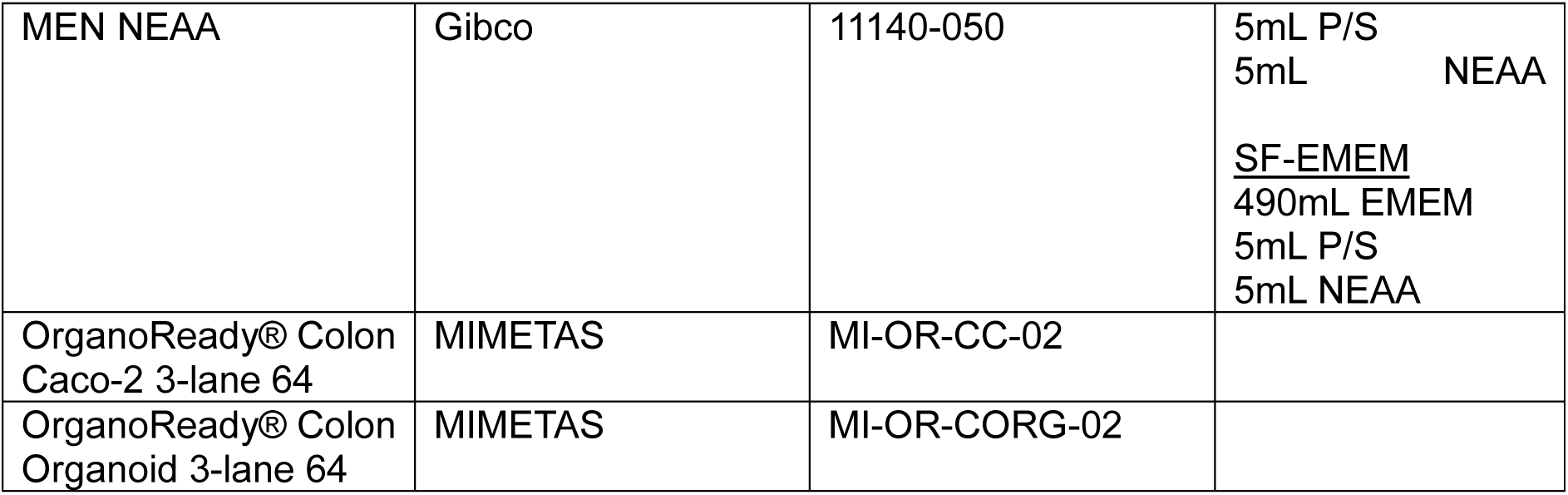

## Supplementary Data

A supplementary dataset is provided to support reproducibility of the protocol. This includes:

- Raw microscopy images used for ACTIN, DRAQ7, and NucBlue staining
- CellProfiler project files for ACTIN and DRAQ7 quantification
- Example output files generated by each pipeline
- A detailed README file explaining the folder structure and instructions to reproduce the analysis

Due to file size limitations, the dataset includes representative raw images from control and treated conditions, along with the CellProfiler pipelines and example output. These materials allow users to replicate the full workflow on a smaller scale, adapt it to their own experimental settings, and validate image analysis outputs. The full dataset (32 chips) is available upon request.

## Acknowledgments

The authors would like to thank their colleagues at MIMETAS and in the FunHoMic consortium for the many fruitful discussions.

## References

1. Rather IA, Koh WY, Paek WK, Lim J (2017) The Sources of Chemical Contaminants in Food and Their Health Implications. Front Pharmacol 8:830. 10.3389/fphar.2017.00830

2. Di Tommaso N, Gasbarrini A, Ponziani FR (2021) Intestinal Barrier in Human Health and Disease. Int J Environ Res Public Health 18:12836. 10.3390/ijerph182312836

3. Eisenbrand G, Pool-Zobel B, Baker V, et al (2002) Methods of in vitro toxicology. Food Chem Toxicol 40:193–236. 10.1016/s0278-6915(01)00118-1

4. Markus J, Landry T, Stevens Z, et al (2021) Human small intestinal organotypic culture model for drug permeation, inflammation, and toxicity assays. In Vitro Cell Dev Biol Anim 57:160–173. 10.1007/s11626-020-00526-6

5. Madorran E, Stožer A, Bevc S, Maver U (2020) In vitro toxicity model: Upgrades to bridge the gap between preclinical and clinical research. Bosn J Basic Med Sci 20:157–168. 10.17305/bjbms.2019.4378

6. Trietsch SJ, Naumovska E, Kurek D, et al (2017) Membrane-free culture and real-time barrier integrity assessment of perfused intestinal epithelium tubes. Nature Communications 8:1–7. 10.1038/s41467-017-00259-3

7. Beaurivage C, Naumovska E, Chang YX, et al (2019) Development of a gut-on-a-chip model for high throughput disease modeling and drug discovery. International Journal of Molecular Sciences 20:. 10.3390/ijms20225661

8. Hagiwara Y, Kumagai H, Ouwerkerk N, et al (2021) A Novel In Vitro Membrane Permeability Methodology Using Three-dimensional Caco-2 Tubules in a Microphysiological System Which Better Mimics In Vivo Physiological Conditions. Journal of Pharmaceutical Sciences 111:214–224. 10.1016/j.xphs.2021.11.016

9. Pöschl F, Höher T, Pirklbauer S, et al (2023) Dose and route dependent effects of the mycotoxin deoxynivalenol in a 3D gut-on-a-chip model with flow. Toxicology in Vitro 105563. 10.1016/j.tiv.2023.105563

10. Morelli M, Cabezuelo Rodríguez M, Queiroz K (2024) A high-throughput gut-on-chip platform to study the epithelial responses to enterotoxins. Sci Rep 14:5797. 10.1038/s41598-024-56520-5

11. Morelli M, Queiroz K (2025) Breaking Barriers: Candidalysin Disrupts Epithelial Integrity and Induces Inflammation in a Gut-on-Chip Model. Toxins 17:89. 10.3390/toxins17020089

12. Lechuga S, Ivanov AI (2021) Actin cytoskeleton dynamics during mucosal inflammation: a view from broken epithelial barriers. Curr Opin Physiol 19:10–16. 10.1016/j.cophys.2020.06.012

13. Heisler DB, Kudryashova E, Hitt R, et al (2024) Antagonistic effects of actin-specific toxins on Salmonella Typhimurium invasion into mammalian cells. bioRxiv 2024.07.01.601609. 10.1101/2024.07.01.601609

