## Supplementary dataset for "Scalable Image-Based Quantification of Cell Permeability and Actin Remodeling: An Example in a Gut-on-Chip Platform": README.txt

Quantification of ACTIN and DRAQ7 Staining in Cytokine-Treated Cells (OrganoPlate 3L64)

This dataset accompanies the protocol used to quantify cytoskeletal integrity (ACTIN) and cell death (DRAQ7) across different cytokine concentrations in a 3-lane OrganoPlate (3L64) using CellProfiler version 4.2.6. Due to file size limitations, only two representative chip are included as example input. The full dataset is available upon request.


Experimental setup
Each chip was treated with a different concentration of cytokines:
E - negative control (vehicle)
N - positive control (cytokines)

Each chip was imaged in three fluorescence channels:
w1 - Hoechst
w2 - DRAQ7
w3 - ACTIN

Folder structure
Dataset/
│
├── Raw_Images/
│   └── [All original microscopy images: Hoechst, DRAQ7, ACTIN]
│
├── ACTIN_ANALYSIS/
│   ├── ACTIN_SCRIPT.cppipe
│   └── ACTIN_OUTPUT/         [CellProfiler output for ACTIN pipeline]
│
└── DRAQ7_ANALYSIS/
    ├── DRAQ7_SCRIPT.cppipe
    └── DRAQ7_OUTPUT/         [CellProfiler output for DRAQ7 pipeline]


Running the Pipelines
To run the pipelines on your system:
1) Open the CellProfiler pipeline file (.cppipe).
2) Navigate to File > Preferences.
Set the appropriate Default Input Folder (folder with raw images to be analyzed) and Default Output Folder based on your local directory structure.

NOTE: The ACTIN_OUTPUT and DRAQ7_OUTPUT folders already contain example output data. Running the pipelines again with the same output path will overwrite these files.
