## Supplementary figures and images for "Scalable Image-Based Quantification of Cell Permeability and Actin Remodeling: An Example in a Gut-on-Chip Platform"

### 130-226_E02_w3B48D5E0A-9B41-4F3D-B8A1-DF6B5DC39CE8.png

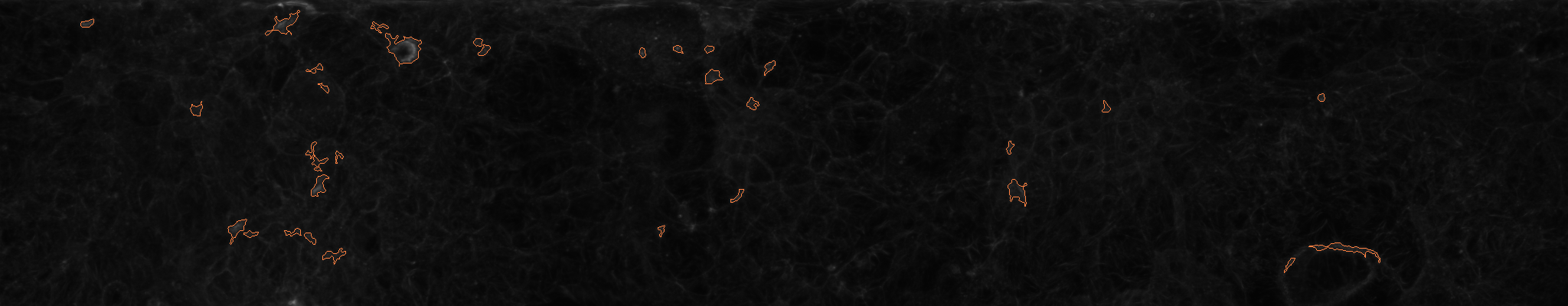

### 130-226_E02_w3B48D5E0A-9B41-4F3D-B8A1-DF6B5DC39CE8.png

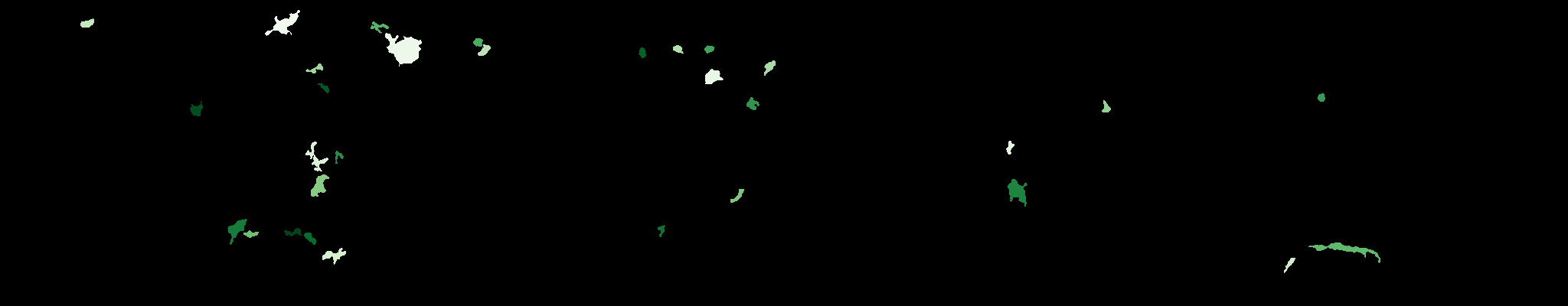

### 130-226_E02_w3B48D5E0A-9B41-4F3D-B8A1-DF6B5DC39CE8.tif

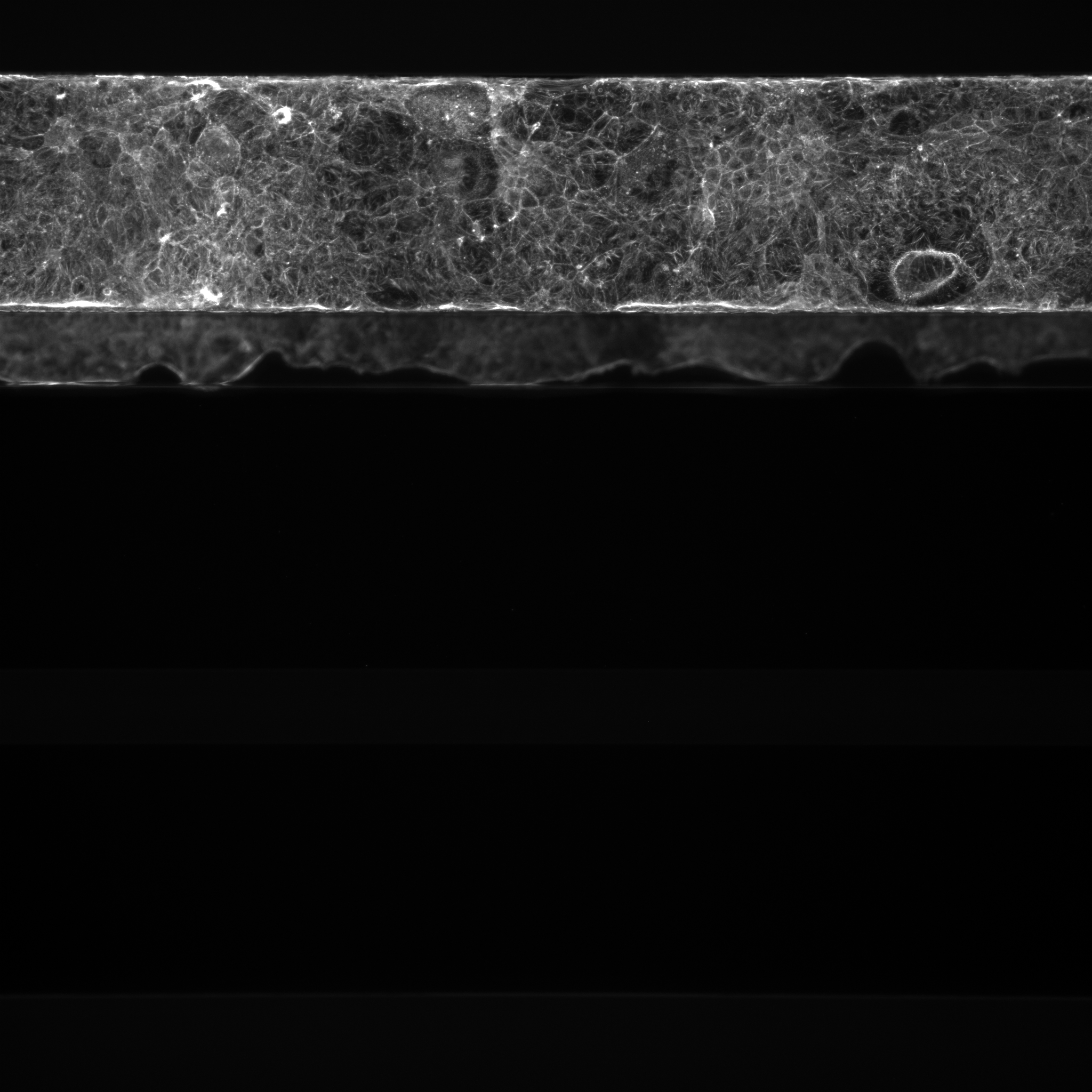

### 130-226_E02_w280050FC3-A3DE-4CE8-A77B-A292308A64B8.png

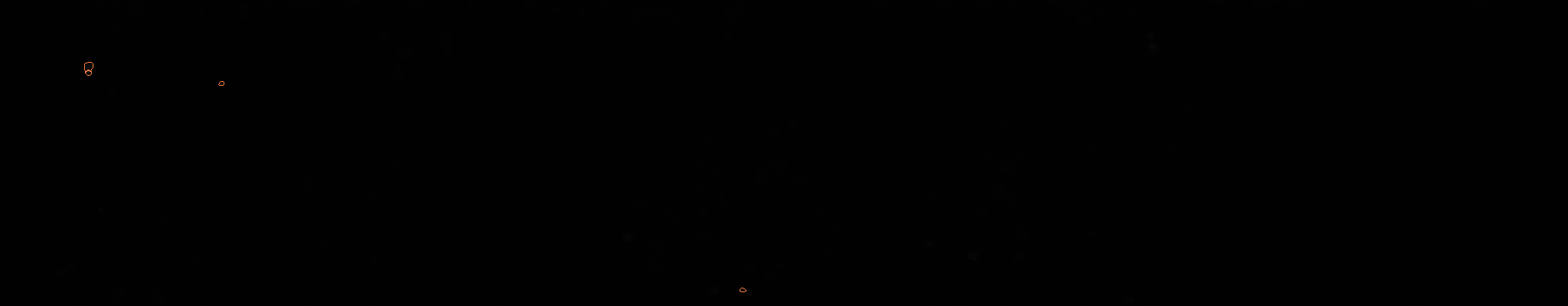

### 130-226_E02_w280050FC3-A3DE-4CE8-A77B-A292308A64B8.png

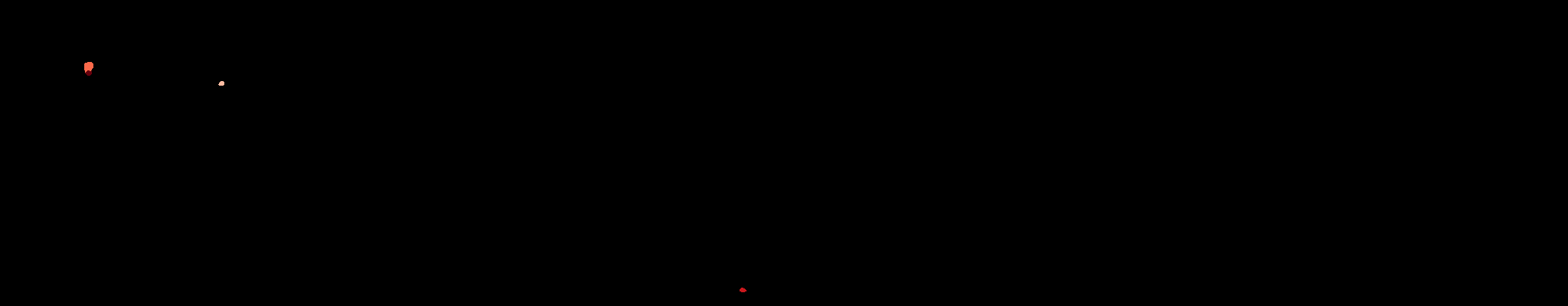

### 130-226_E02_w280050FC3-A3DE-4CE8-A77B-A292308A64B8.tif

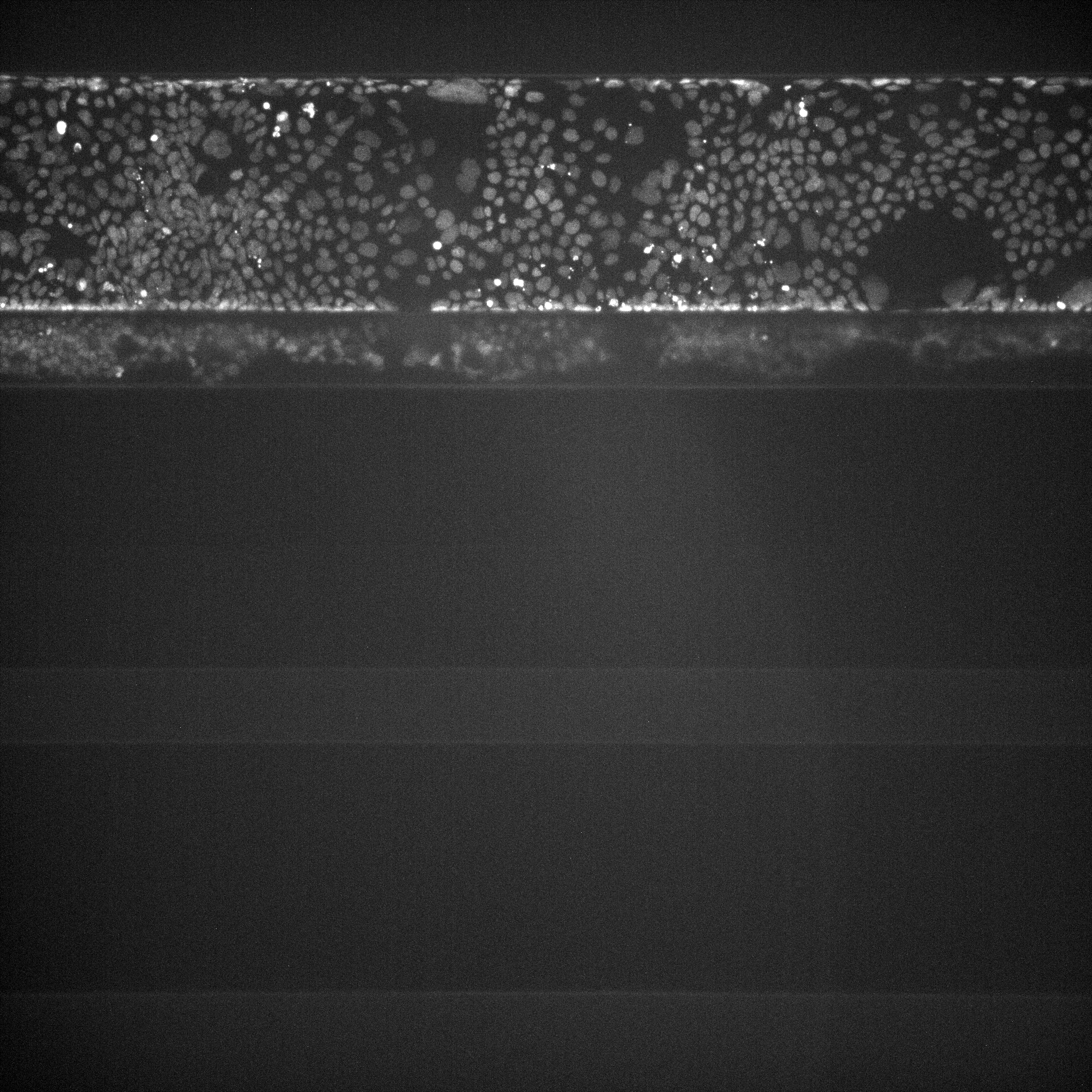

### 130-226_E02_w123497361-86CE-4A69-8E90-27B97F52779A.png

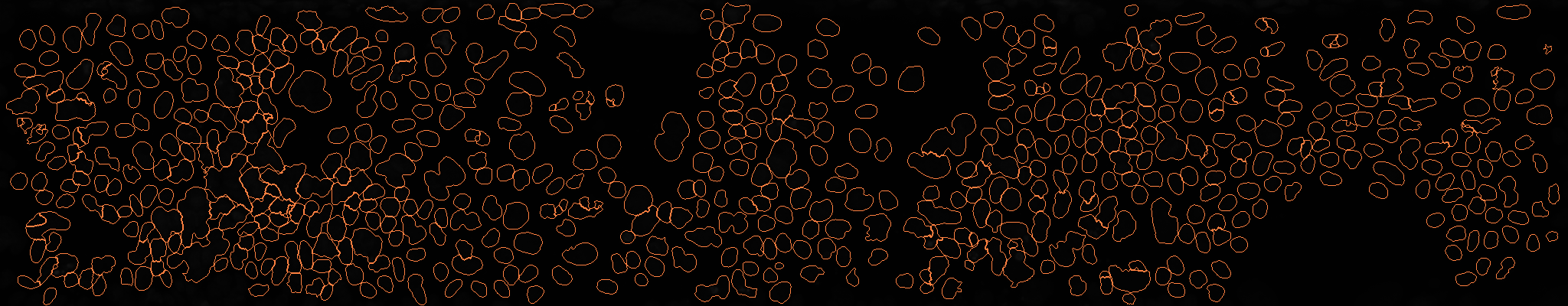

### 130-226_E02_w123497361-86CE-4A69-8E90-27B97F52779A.png

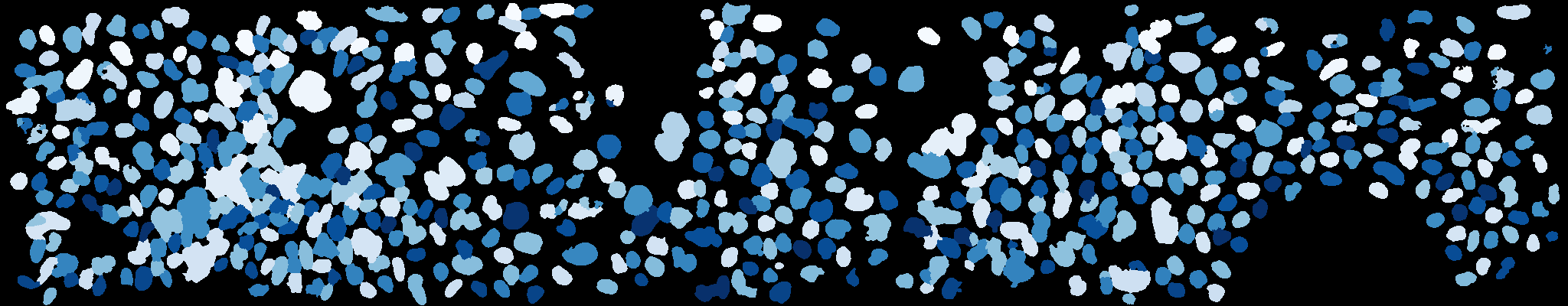

### 130-226_E02_w123497361-86CE-4A69-8E90-27B97F52779A.tif

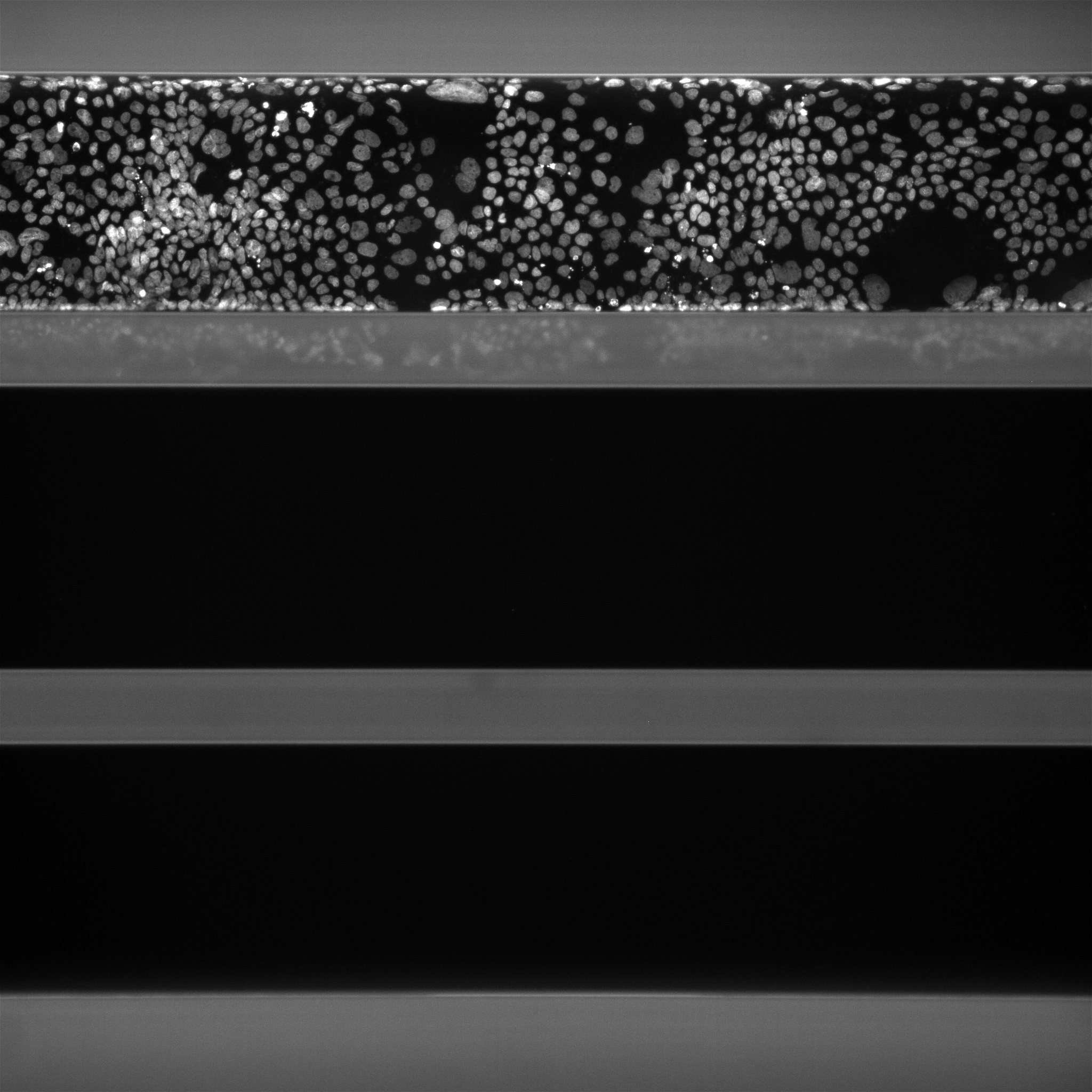

### 130-226_N08_w3EFDA1C76-5F85-47F1-9FCA-324DF0A77F72.png

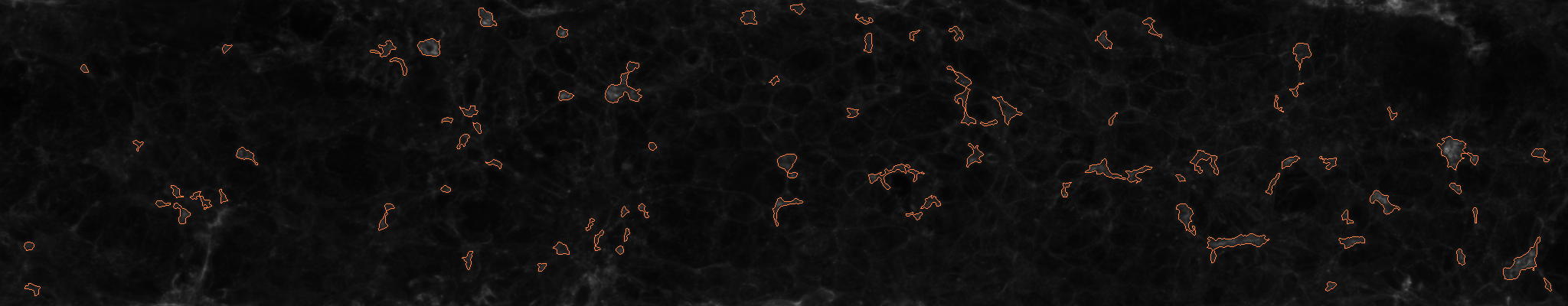

### 130-226_N08_w3EFDA1C76-5F85-47F1-9FCA-324DF0A77F72.png

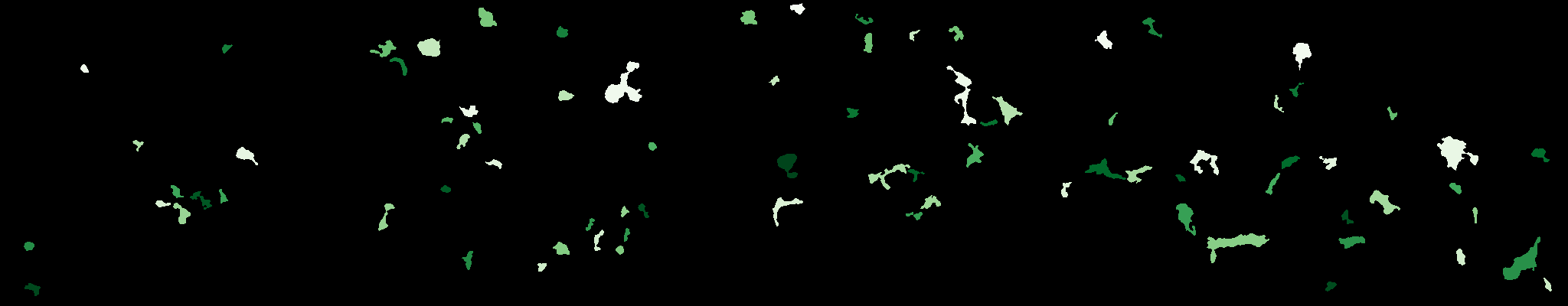

### 130-226_N08_w3EFDA1C76-5F85-47F1-9FCA-324DF0A77F72.tif

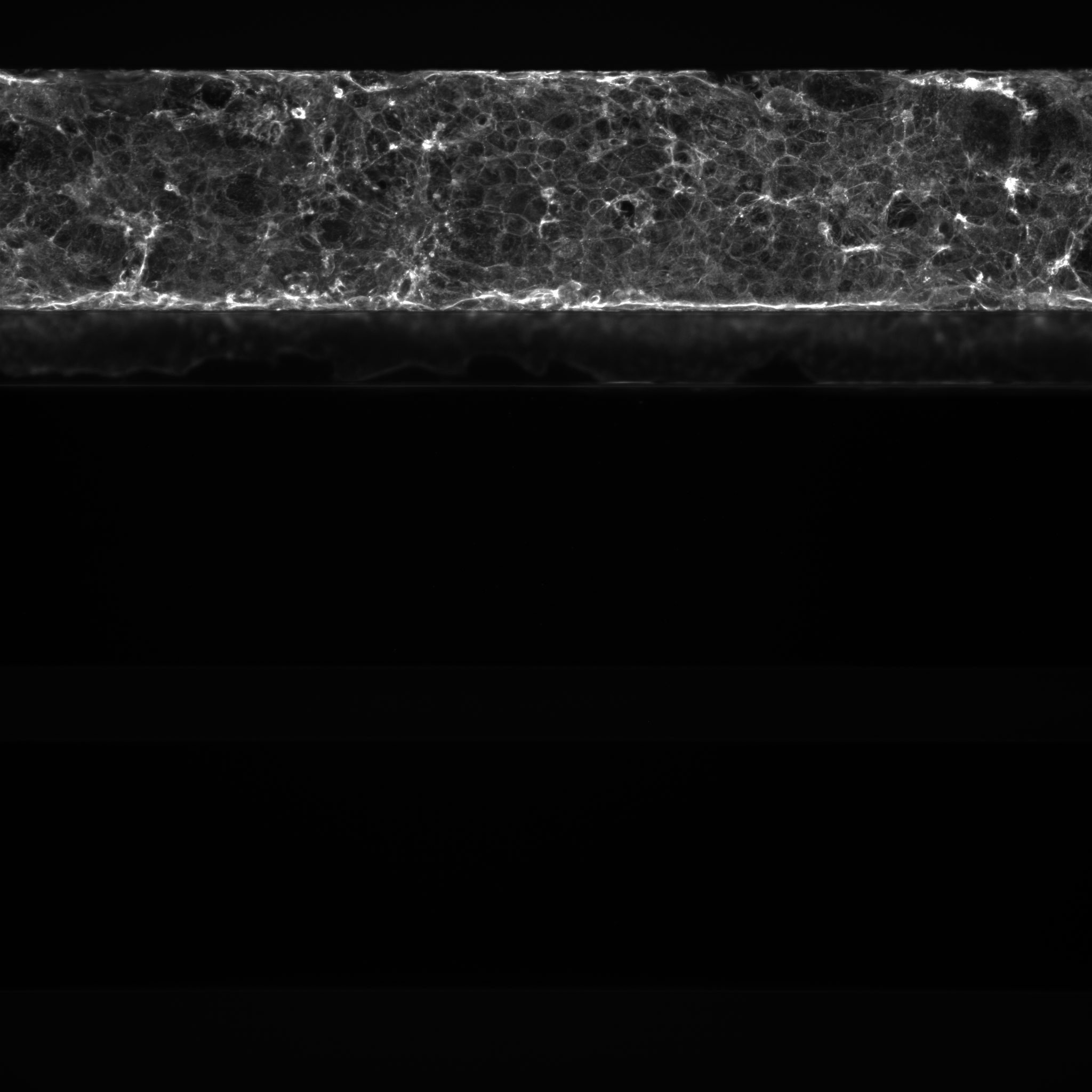

### 130-226_N08_w103CF5AF1-103E-4C28-A723-62016FD9693D.png

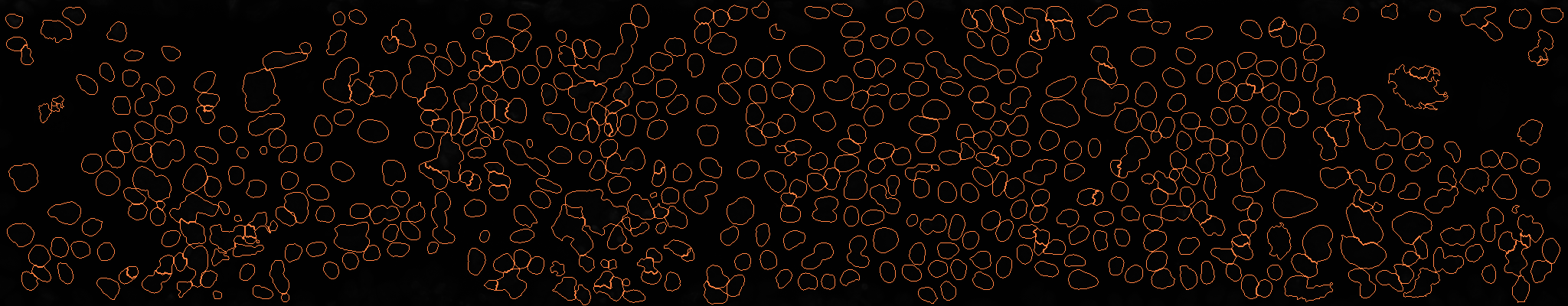

### 130-226_N08_w103CF5AF1-103E-4C28-A723-62016FD9693D.png

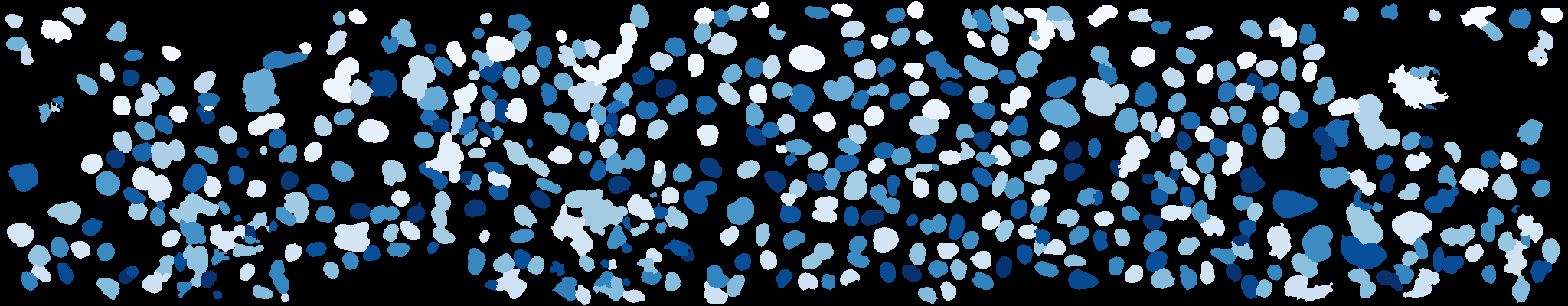

### 130-226_N08_w103CF5AF1-103E-4C28-A723-62016FD9693D.tif

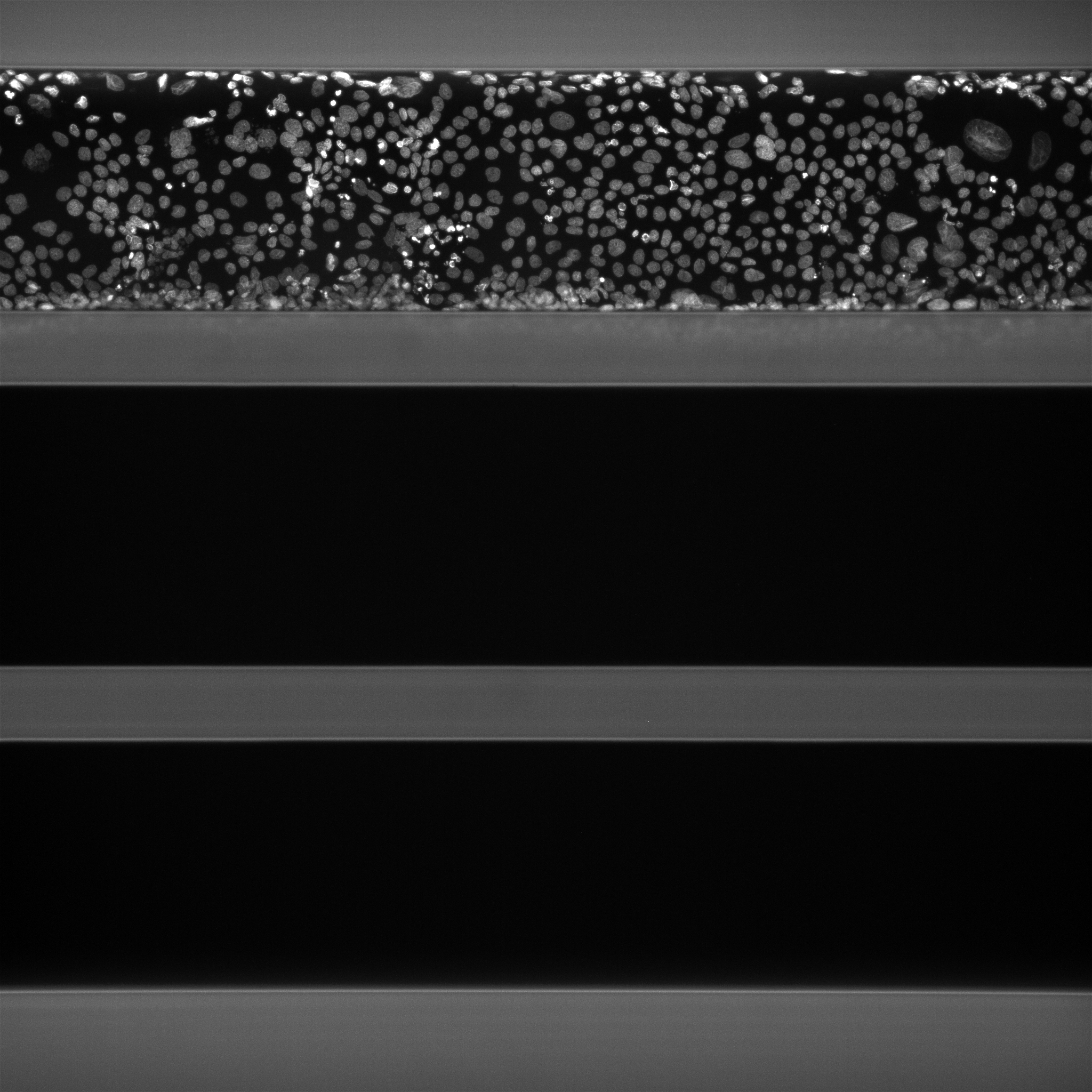

### 130-226_N08_w2503AB1FC-1907-4DF9-A5DD-B1BD9D777BF0.png

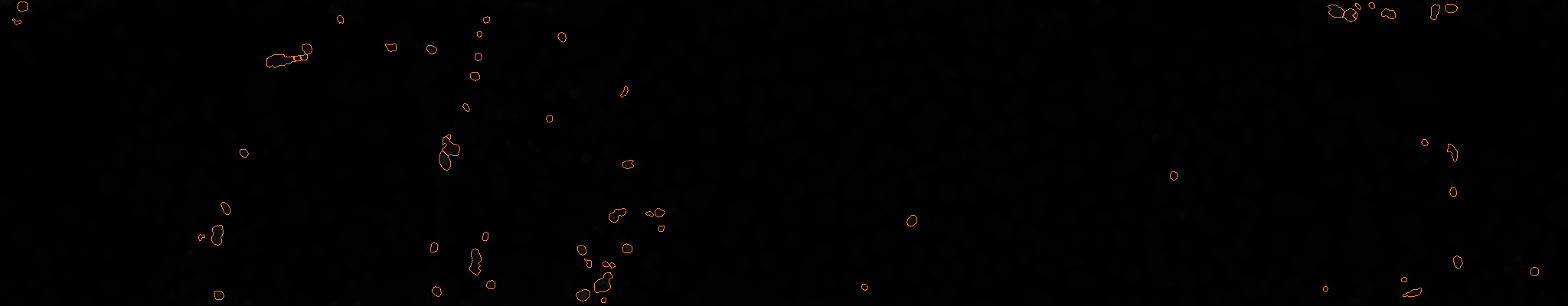

### 130-226_N08_w2503AB1FC-1907-4DF9-A5DD-B1BD9D777BF0.png

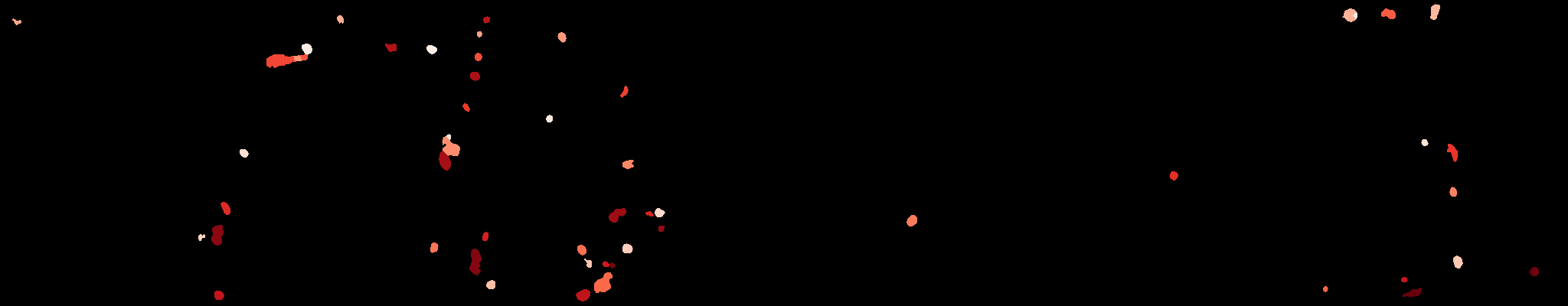

### 130-226_N08_w2503AB1FC-1907-4DF9-A5DD-B1BD9D777BF0.tif

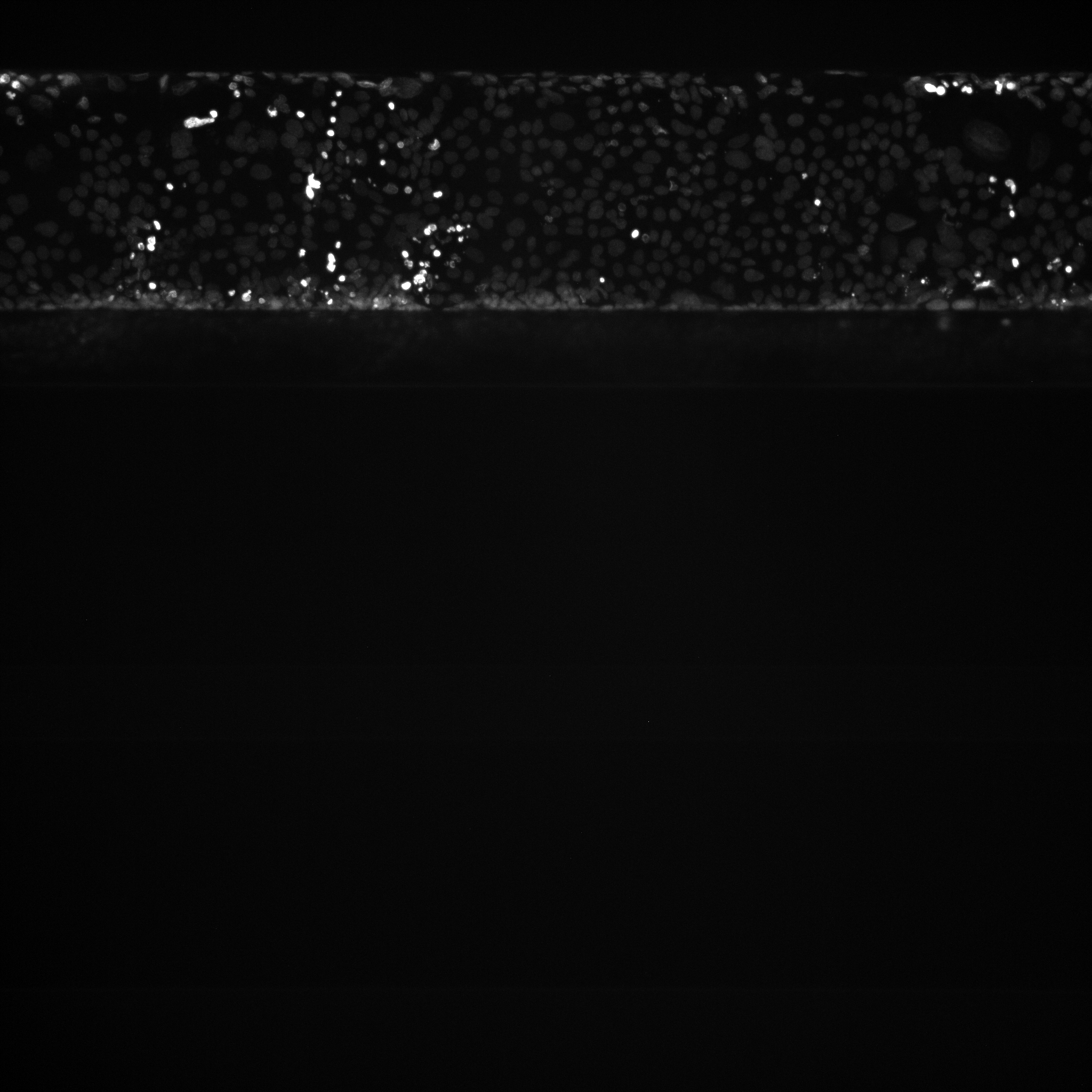
